# Genotype-immunophenotype relationships in *NPM1*-mutant AML clonal evolution uncovered by single cell multiomic analysis

**DOI:** 10.1101/2024.11.11.623033

**Authors:** Morgan Drucker, Darren Lee, Xuan Zhang, Bailee Kain, Michael Bowman, Deedra Nicolet, Zhe Wang, Richard M. Stone, Krzysztof Mrózek, Andrew J. Carroll, Daniel T. Starczynowski, Ross L. Levine, John C. Byrd, Ann-Kathrin Eisfeld, Nathan Salomonis, H. Leighton Grimes, Robert L. Bowman, Linde A. Miles

**Affiliations:** Division of Hematology/Oncology, Cancer & Blood Disease Institute, Cincinnati Children’s Hospital Medical Center, Cincinnati OH USA; University of Cincinnati College of Medicine, Cincinnati OH USA; Division of Immunobiology, Cincinnati Children’s Hospital Medical Center, Cincinnati OH USA; Department of Cancer Biology, Perelman School of Medicine, University of Pennsylvania, Philadelphia PA USA; The Ohio State University Comprehensive Cancer Center, Columbus, OH USA; Clara D. Bloomfield Center for Leukemia Outcomes Research, The Ohio State University Comprehensive Cancer Center, Columbus OH USA; Dana-Farber/Partner CancerCare, Boston MA, USA; Department of Genetics, University of Alabama at Birmingham, Birmingham, AL USA; Division of Experimental Hematology & Cancer Biology, Cancer & Blood Diseases Institute, Cincinnati Children’s Hospital Medical Center, Cincinnati OH USA; Department of Pediatrics, University of Cincinnati, Cincinnati OH USA; University of Cincinnati Cancer Center, Cincinnati OH USA; Human Oncology and Pathogenesis Program, Molecular Cancer Medicine Service, Memorial Sloan Kettering Cancer Center, New York, New York, USA; Center for Hematologic Malignancies, Memorial Sloan Kettering Cancer Center, New York, NY, USA; Leukemia Service, Department of Medicine, Memorial Sloan Kettering Cancer Center, New York, NY, USA; Department of Internal Medicine, University of Cincinnati, Cincinnati OH USA; Division of Hematology Department of Internal Medicine, The Ohio State University Comprehensive Cancer Center, Columbus, OH USA; Division of Biomedical Informatics, Cincinnati Children’s Hospital Medical Center, Cincinnati, OH USA

## Abstract

Acute myeloid leukemia (AML) is a multi-clonal disease, existing as a milieu of clones with unique but related genotypes as initiating clones acquire subsequent mutations. However, bulk sequencing cannot fully capture AML clonal architecture or the clonal evolution that occurs as patients undergo therapy. To interrogate clonal evolution, we performed simultaneous single cell molecular profiling and immunophenotyping on 43 samples from 32 *NPM1*-mutant AML patients at different stages of disease. Here we show that diagnosis and relapsed AML samples display similar clonal architecture patterns, but signaling mutations can drive increased clonal diversity specifically at relapse. We uncovered unique genotype-immunophenotype relationships regardless of disease state, suggesting leukemic lineage trajectories can be hard-wired by the mutations present. Analysis of longitudinal samples from patients on therapy identified dynamic clonal, transcriptomic, and immunophenotypic changes. Our studies provide resolved understanding of leukemic clonal evolution and the relationships between genotype and cell state in leukemia biology.

## Main

Acute myeloid leukemia (AML) is an aggressive blood cancer that arises from the aberrant expansion of mutant hematopoietic stem and progenitor cells, which leads to the blockade of normal differentiation. Variant allele frequencies (VAF) inferred from large-scale bulk sequencing studies largely suggest that AML initiating mutations in epigenetic regulators (*TET2, DNMT3A, IDH1/2*) are followed by mutations in signaling genes (*RAS, FLT3*)^1-3^. One of the most recurrently mutated AML genes is nucleophosmin 1, *NPM1*, which is mutated in approximately 30% of adults with AML^1-3^. *NPM1*-mutant AML is considered a distinct disease entity by both the World Health Organization (WHO) and International Consensus Classification (ICC)^4,5^ and typically harbors epigenetic modifier and/or signaling gene co-mutations*^6,7^*. However, the synergistic interactions of these mutations and their contributions towards clonal fitness and transformation remain to be uncovered.

Recent large cohort single cell multiomic (DNA + cell surface protein expression) studies by us and others have assessed the clonal architecture of myeloid malignancies, including AML, and provided improved resolution to AML clonal heterogeneity^8-11^. These studies revealed that mutations in epigenetic regulators vs signaling genes have different representation in the dominant clone, and mutational combinations may affect lineage output^8,9^. Longitudinal sampling of AML patients while undergoing targeted therapy with FLT3 or IDH inhibitors were performed in small cohorts and suggested significant dynamics in clones over time and while under selective pressure^12-14^. These bulk sequencing and single cell multiomic studies highlight the need to better understand clonal evolution while patients undergo therapy and to assess how specific combinations of mutations may create divergent evolutionary trajectories for leukemia even within similarly classified AML patients (i.e. *NPM1*-mutant AML).

In this study, we perform single cell multiomic analysis on 609,314 cells in 43 samples from 32 *NPM1*-mutated AML patients to interrogate how different co-mutations may dictate evolutionary trajectories for mutant clones. We first interrogate clonal architecture patterns in *NPM1*-mutated AML across different disease states and how co-mutations affect clonal framework patterns in individual patient samples. We identify genotype-immunophenotype correlations within the cohort to understand how co-mutations affect differentiation patterns in AML. We next analyze clonal evolution in longitudinal samples from 8 patients who underwent 7+3 chemotherapy and identify distinct patterns in clonal changes, even across patients with the same genotype. Using a complementary single cell multiomic approach, CITE-seq, we further investigate gene expression differences at diagnosis and relapse, unveiling significant alterations in connected signaling cascades and protein ubiquitination pathways, suggestive of alternative signaling as cells respond to therapy.

## Results

### Clonal architecture patterns suggest similar heterogeneity levels between diagnosis and relapse samples

We performed simultaneous single cell molecular profiling and cell surface protein expression (DNA+Protein) sequencing on 609,314 cells from 43 samples from 32 patients with *NPM1*-mutated AML. *NPM1* and all co-mutations were initially identified and confirmed through targeted bulk sequencing (**Fig. 1ab**; **Extended Table 1**). The most common co-mutations identified with bulk sequencing were in *FLT3* (n = 17), *IDH1/2* (n = 15), and *TET2 (*n = 14). Eighty seven percent of patients had two or more mutations in addition to *NPM1* mutations. We queried samples from patients at different stages of disease, including diagnosis (n = 20), complete response (CR; n = 4) while on therapy, and relapse (n = 19) (**Fig. 1b; Extended Data Fig. 1**). For 24 patients, we sequenced a single sample from their disease course. Eight patients from our cohort were longitudinally sampled (2-3 samples) while on variations of a standard cytotoxic chemotherapy regimen, known as 7+3^15^, which consists of 7 days of continuous cytarabine with 3 days of dauno/doxo-rubicin (**Extended Data Fig. 1; Extended Table 2**). For each patient we generated a clonograph to determine the abundance and heterogeneity of clones present in each patient (**Fig. 1c**).

**Fig. 1.**
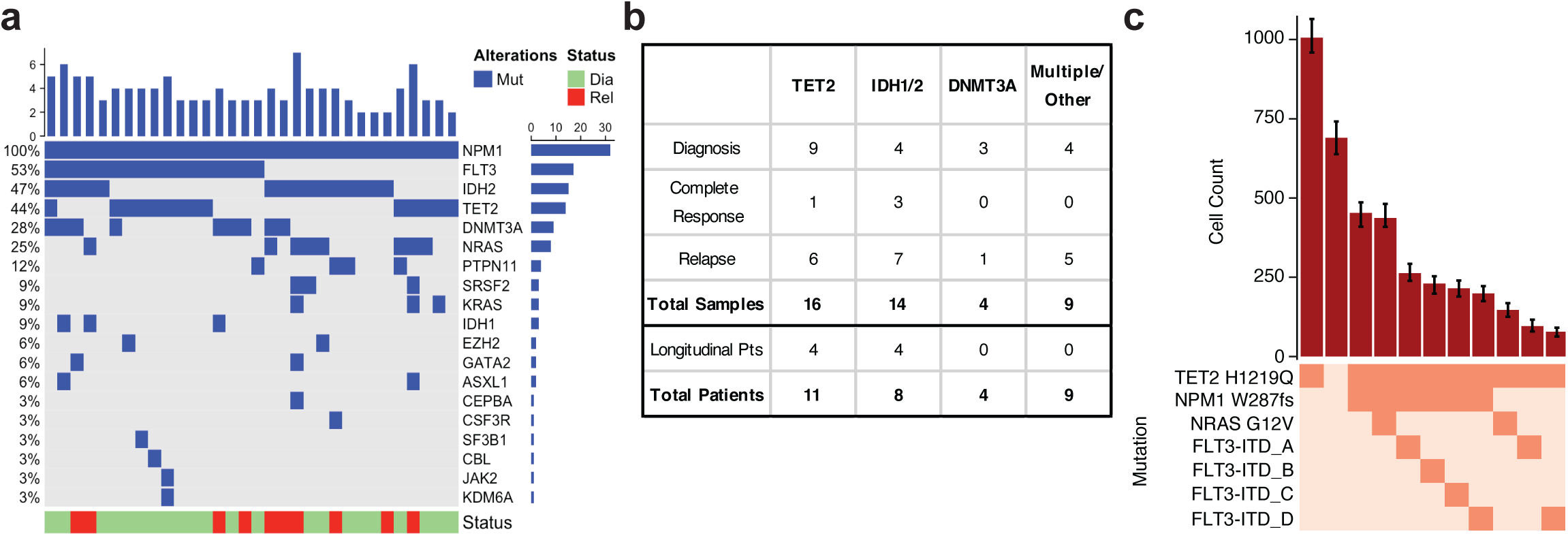
*NPM1*-mutated AML patient cohort. **a)** Oncoprint of samples in patient cohort (n=32) depicting mutations identified by targeted bulk sequencing. For patients with more than one sample, only the diagnosis sample is displayed. **b)** Table of patient cohort (n=43 samples) describing breakdown of samples by epigenetic co-mutation and disease state. **c)** Clonograph of a representative patient sample (Pt I diagnosis) depicting clones present in sample. The height of each bar represents the cell count of the clone identified below. Clone genotype is depicted by color with wildtype (WT) in light beige and heterozygous mutations in orange denoted.

We first investigated differences in clonal architecture between the various disease states: diagnosis, CR, and relapse across the entire cohort. There were no significant differences in the number of mutations per sample or the number of mutations in the dominant clone (defined as the largest non-wildtype clone) between disease states (**Extended Data Fig. 2ab**), suggesting that alterations to mutational burden are not the main driver of response or relapse. No significant difference in dominant clone size between diagnosis and relapse was observed (**Fig. 2a**). There was, however, a significant decrease in the number of distinct clones per sample (*P* = 0.007) and Shannon diversity index (*P* = 0.005) from diagnosis to CR and subsequent increase in these same parameters from CR to relapse (number of clones, *P* = 0.03; Shannon diversity index, *P* = 0.03; **Fig. 2bc**). This pattern suggests that the clonal heterogeneity observed at initial diagnosis returns with relapse through expansion of the existing clones and/or the development of new clones that replace ones lost during therapy and response.

**Fig. 2.**
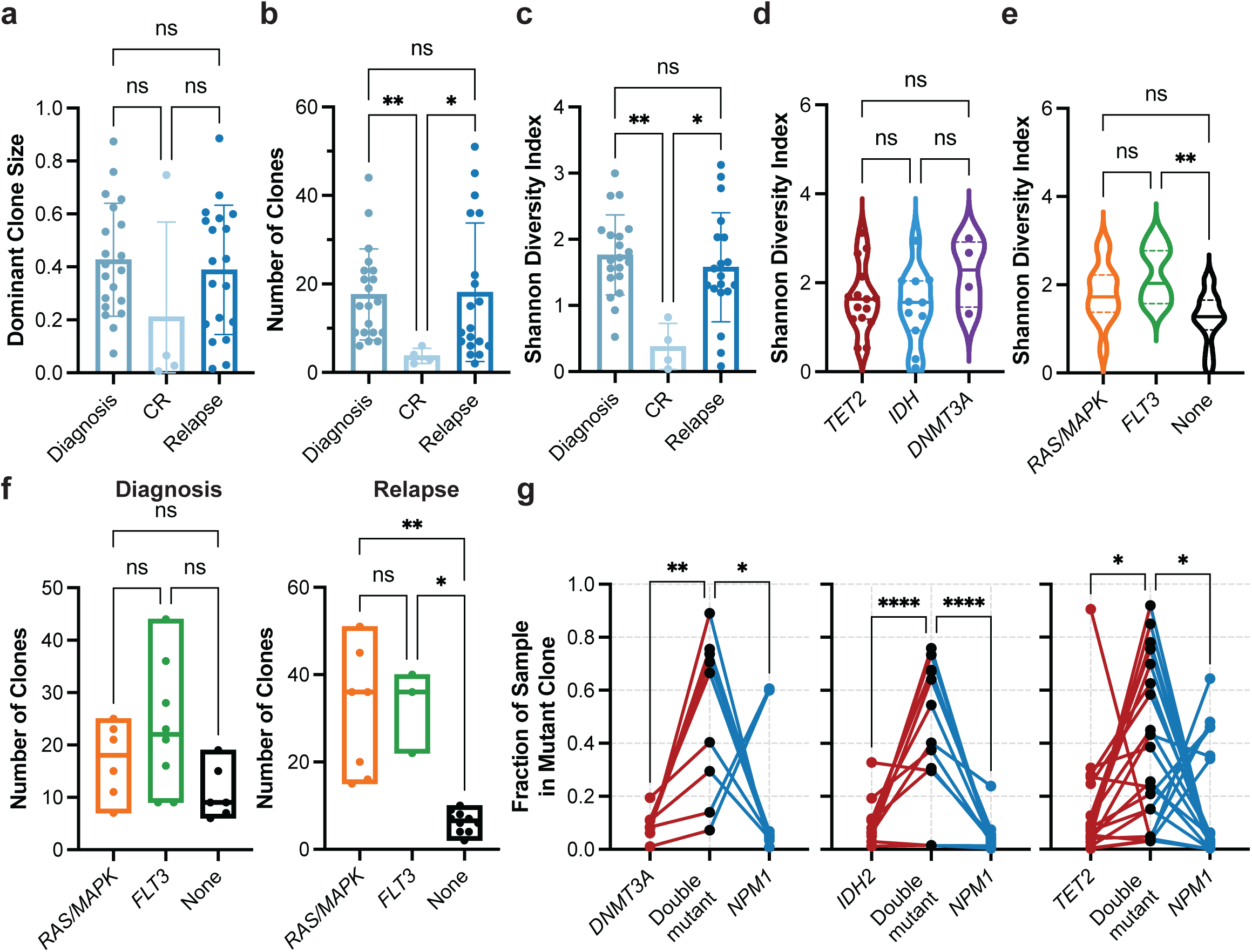
Clonal architecture patterns and mutational cooperativity by single-cell DNA sequencing. **a-c)** Bar graphs of clonal architecture metrics for entire cohort by disease state, including (**a**) dominant clone size, (**b**) number of clones, and (**c**) clonal diversity, calculated by the Shannon diversity index (mean ± SD, n = 43). **d)** Violin plot of Shannon diversity index for diagnosis and relapse samples harboring epigenetic mutations (n = 30) in *TET2* (red), *IDH1/2* (blue) or *DNMT3A* (purple). Samples with more than one epigenetic mutation were excluded from analysis. **e)** Violin plot of Shannon diversity index for diagnosis and relapse samples with *RAS/MAPK* (n=13; orange), *FLT3* (n=11; green), or no signaling gene mutations (n=16; None; black). Samples with both a RAS/MAPK and FLT3 mutation were excluded. **f)** Number of clones identified in samples with *RAS/MAPK* (n=13; orange), *FLT3* (n=11; green), or no signaling gene mutations (n=16; None; black) stratified by disease state (Diagnosis, left panel; Relapse, right panel). Kruskal-Wallis test was used to determine statistical significance amongst groups (**a-f**). **g)** Fraction of sample in single- and double-mutant clones in samples with *DNMT3A-NPM1* (n = 9; left panel), *IDH2-NPM1* (n = 12; center panel), and *TET2-NPM1* (n = 20; right panel) mutations. Individual samples denoted by connecting lines. Two-way ANOVA used to determine statistical significance (**g**) *P<0.05, **P<0.01, ***P<0.001 denoted for all panels.

### Presence of signaling mutations correlate with increased heterogeneity at relapse

We then examined whether there were differences in the clonal framework among diagnosis and relapse samples classified by co-mutations in epigenetic modifier genes (*IDH1/2, TET2, DNMT3A)* versus stratification based on the presence or absence of signaling gene mutations (*FLT3, RAS/MAPK*). We observed no differences in the number of mutations/clones or dominant clone size between samples harboring different epigenetic gene mutations (**Extended Data Fig. 2cde**). Correspondingly, we also did not see significant differences in clonal diversity between samples stratified by epigenetic gene mutations (**Fig. 2d**). However, samples with mutations in signaling genes, *RAS/MAPK* and *FLT3*, were observed to have an increased number of mutations (None vs *RAS P* = 0.004, vs. *FLT3 P* = 0.0003) and clones (None vs *RAS P* = 0.0003, vs *FLT3 P* = 0.0002) compared to samples without signaling mutations with no notable difference in the dominant clone size (**Extended Data Fig. 2fgh**). Further, *FLT3* mutant samples were found to have increased clonal diversity as compared with samples without any signaling mutations (*P* = 0.002; **Fig. 2e**). Critically, this signaling mutant-driven increase in clonal complexity was uniquely identified in the relapse setting, as we did not observe significant differences in the clonal metrics in diagnosis samples (**Extended Data Fig. 2ij**, **Fig. 2f**). These findings significantly improve the resolution and add clinical context for a similar pattern we observed previously in a larger cohort identifying an increase in clonal diversity in AML samples with mutations in signaling genes^8^. Moreover, these findings suggest that patients who harbor signaling gene mutations may undergo relapse through increasing overall clonal diversity and polyclonality, where multiple clones are competing for increased clonal fitness and dominance. However, patients without signaling gene mutations do not show increased clonal heterogeneity in the relapse setting, suggesting they may utilize alternative mechanisms to drive relapse.

### Mutational cooperativity levels vary based on co-mutation

We next investigated patterns of mutational cooperativity in *NPM1*-mutated AML samples. Our previous study suggested that *NPM1* mutations may drive clonal expansion when co-mutant with epigenetic modifiers and signaling mutations, albeit to varying degrees based on the co-mutation^8^. Aligning with our previous study, we observed similar patterns across all epigenetic co-mutations, with an increased relative clone size of double-mutant clones compared with single-mutant *TET2* (*P* = 0.03)*, IDH2* (*P* < 0.0001), or *DNMT3A* clones (*P* = 0.01) and/or single-mutant *NPM1* clones (*TET2 P* = 0.03*; IDH2 P* < 0.0001*; DNMT3A P* = 0.03) identified in the sample (**Fig. 2g**). Between the epigenetic mutations, we noted a stronger trend towards increased double mutant clone size for *IDH2/NPM1* co-mutant clones compared to *DNMT3A* or *TET2* co-mutant clones. Conversely, for samples with co-occurring signaling gene mutations, there was more variability in the size of *NPM1* single mutant clones with less evidence of cooperativity in the *NPM1-RAS* or *NPM1-FLT3* double mutant clones (**Extended Data Fig. 2k**). Significant clonal expansion was more evident when comparing single mutant *FLT3* (*P* = 0.003) or *NRAS* (*P* = 0.04) to double mutant clones. These studies suggest that mutational cooperativity is highly context dependent and may vary significantly based on co-mutation identity and the synergy between the cellular alterations imparted by both *NPM1* and the co-mutation.

### Immunophenotypic analysis reveals lineage biases across disease stages

We next assessed cell surface protein expression across the sample cohort at single cell resolution. Analysis of the single cell surface protein expression (scProtein) confirmed that overlapping immunophenotypes could be observed across individual samples and patients within the cohort (**Extended Data Fig. 3a**). Cells were then clustered into 31 unique communities based on similarities in their aggregated expression of measured cell surface markers with each community defined by the expression of more than one marker (**Fig. 3a; Extended Data Fig. 3bc**). Upon stratification of samples based on disease stage (Diagnosis, CR, Relapse), we found that the heterogeneity of community representation calculated by a Shannon index is not significantly different (**Extended Data Fig. 3de**). However, we did observe certain communities that were enriched or depleted in representation based on disease stage (**Fig. 3b**, **Extended Data Fig. 3d**). CR samples were enriched in representation in clusters 3 and 4, which contained 28.7% (cluster 3 = 16.5%; cluster 4 = 12.2%) of total cells from CR samples, compared to 8.8% (cluster 3 = 3.8%; cluster 4 = 5.0%) and 7.1% (cluster 3 = 3.5%; cluster 4 = 3.6%) of cells from Diagnosis and Relapse samples, respectively. Clusters 3 and 4 harbored cells with the highest expression of the classical T-cell markers CD3, CD4, and CD8 (**Fig. 3bc, Extended Data Fig. 3c**). Leukemic samples (Diagnosis/Relapse), on the other hand, were found to be enriched in clusters 0, 2, and 16 which contained 21.8% and 25.9% of total cells from Diagnosis and Relapse samples, respectively. These clusters expressed higher levels of CD38, CD117 and CD123, known markers for leukemic blasts as well as enrichment of CD11b, CD64, and CD14 shown to be expressed on monocytic AML blasts and myeloid progenitors (**Fig. 3bc, Extended Data Fig. 3c**). There were certain clusters with stable representation regardless of disease stage, including clusters 6, enriched in B-cell markers CD19 and CD22 (3.8% Diagnosis, 3.9% CR, 4.2% Relapse). Lastly, we observed similar representation in cluster 8 across disease states (3.4% Diagnosis, 3.0% CR, 3.2% Relapse), which express CD11b, CD64, and CD14 without stem/progenitor markers and cluster 11 (3.4% Diagnosis, 3.8% CR, 2.7% Relapse) with expression of promyelocytic markers CD141, CD71 and CD49d, suggesting that certain immunophenotypes are always generated regardless of disease state.

**Fig. 3.**
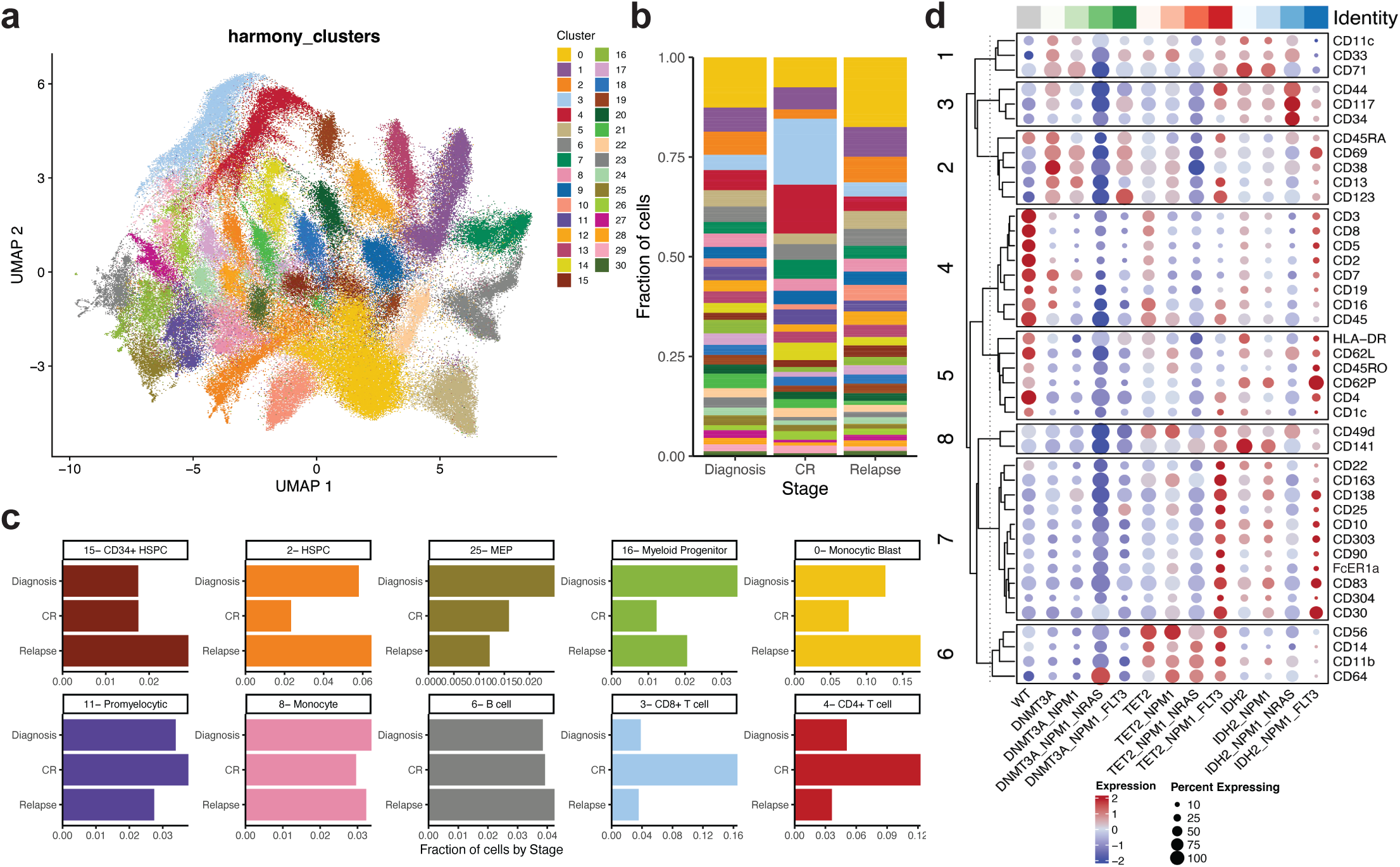
Identification of genotype-immunophenotype relationships using simultaneous single-cell DNA+Protein sequencing. **a)** Uniform manifold approximation and projection (UMAP) plot of 31 communities identified based on aggregate protein data from entire patient cohort (n=43) with cells clustered by immunophenotype. **b)** Fraction of cells within a given disease stage (diagnosis, CR, relapse) clustered into the 31 communities previously identified across cohort with colors matching community identity in Fig. 3a. **c)** Bar graphs depicting fraction of cells from a given disease state (diagnosis, CR, relapse) identified within a community. Community number with corresponding immunophenotype signature based on immunophenotype markers enriched within that community denoted. Colors of community denoted in UMAP in Extended Data Fig. 3b. **d)** Dot plot depicting expression of immunophenotypic markers by genotype-specific clones. Normalized expression of each marker depicted by color (blue = low, red = high) with size of dot denoting the fraction of cells within each genotyped clone that expresses the marker. Immunophenotype markers grouped by corresponding lineage associations. Top bar, gray = WT, green spectrum = *DNMT3A* clones, red spectrum = *TET2* clones, blue spectrum = *IDH2* clones. Full genotype for each column denoted at bottom of the dotplot.

### scProtein uncovers genotype-specific immunophenotypic patterns

We next examined how specific genotypes within *NPM1*-mutant AML affected immunophenotypes and lineage biases. We observed that genotypes could indeed alter lineage output, albeit to varying degrees, aligning with our previous findings with a smaller initial scProtein panel^8^. We observed that all mutant cells were significantly excluded from the T-cell clusters, to different degrees depending upon the mutations. Amongst the mutated cells, *TET2* mutant cells were the most abundant (odds ratio [OR] = 0.379 and 0.292 for clusters 3 and 4, respectively) whilst *NPM1* and *FLT3* were markedly rare (*NPM1* OR = 0.057 and 0.063, *NRAS* OR = 0.159 and 0.157 for clusters 3 and 4, respectively; **Extended Data Fig. 3d**). We next grouped clones by genotype from the entire cohort and investigated alterations to marker expression. We uncovered stark contrasts between clones harboring *DNMT3A, TET2*, and *IDH2* (**Fig. 3d; Extended Data Fig. 3c**). Clones harboring *DNMT3A*-mutations were enriched for higher CD38 expression and lower CD11b expression compared to clones harboring *TET2*- and *IDH2*-mutations (avg scaled expression CD38: *DNMT3A*-clones 0.40+/-1.32 vs *IDH2*-clones 0.21+/-0.23 or *TET2*-clones - 0.42+/-1.22; CD11b: *DNMT3A*-clones -1.24+/-0.30 vs *IDH2*-clones 0.40+/-0.32 or *TET2*-clones 1.00+/-0.26). Clones harboring *IDH2*-mutations showed increased expression of stem/progenitor markers such as CD141 previously suggested to represent neoplastic clones^16^, as well as CD34 and CD117 (avg scaled expression CD141: *IDH2*-clones 0.81+/-0.90 vs *DNMT3A*-clones -1.05+/- 0.76 or *TET2*-clones 0.30+/-0.33; CD34: *IDH2*-clones 0.92+/-1.06 vs *DNMT3A*-clones -0.43+/- 0.52 or *TET2*-clones -0.36+/-0.45; CD117: *IDH2*-clones 0.90+/-0.82 vs *DNMT3A*-clones -0.43+/- 1.05 or *TET2*-clones -0.23+/-0.71). Strikingly, clones harboring *TET2*-mutations diverged significantly from *DNMT3A-* and *IDH2*-mutant clones in that they were instead enriched for markers including CD14, CD11b and CD64, prominent mature monocytic markers (avg scaled expression CD14: *TET2*-clones 1.20+/-0.59 vs *DNMT3A*-clones -0.79+/-0.34 or *IDH2*-clones - 0.42+/-0.73; CD64: *TET2*-clones 0.78+/-0.50 vs *DNMT3A*-clones -0.36+/-1.3 or *IDH2*-clones 0.02+/-0.24). These findings suggest that epigenetic mutations including *DNMT3A*, *TET2*, and *IDH2* may dictate lineage biases and differentiation potential that is inherited by subsequent clones.

Next, we found that *NRAS*- and *FLT3*- mutant clones of the same epigenetic genotype possessed different immunophenotypic patterns (**Fig. 3d; Extended Data Fig. 3c**). Compared to *DNMT3A/NPM1/NRAS*-mutant cells (DNR), *DNMT3A/NPM1/FLT3* co-mutant cells (DNF) expressed 25.1-fold more CD123 and 14.0-fold more CD117, the latter being previously suggested by immunophenotyping studies^17,18^ (average expression CD123: DNF: 55.9 vs DNR: 2.22; CD117: DNF: 29.6 vs DNR: 2.1). *TET2*-mutant clones harbored similar patterns with *TET2/NPM1/FLT3* co-mutant clones showing higher expression of CD117 and CD123 compared to *TET2/NPM1/NRAS* mutant clones (average expression CD117: TNF: 44.1 vs TNR: 7.28; CD123: TNF: 55.5 vs TNR: 6.75). In *IDH2* mutant clones, an opposite trend was observed with *IDH2/NPM1/NRAS* mutant clones expressing 5.4-fold higher CD117 and 7.1-fold higher CD34 compared to *IDH2/NPM1/FLT3* co-mutant clones, suggesting the *IDH2/NPM1/NRAS* combination harbors a strong stem/progenitor phenotype (average expression CD117: INR: 186.6 vs INF: 34.8; CD34: INR: 40.7 vs INF: 5.69). Moreover, *FLT3*-mutant clones showed increased expression of CD25, previously reported as a biomarker of *FLT3*-mutant cells^19^ as well as CD30 and CD69, both of which are known to be expressed on AML blasts with increased self-renewal and stem-like properties^20,21^. These results imply that signaling mutations can refine further lineage trajectories established by epigenetic mutations in antecedent clones, creating unique genotype-immunophenotype relationships.

### Longitudinal sampling of patients during therapy undercovers genotypic and immunophenotypic clonal evolution

To investigate how standard cytotoxic chemotherapy affects patients on a clonal and immunophenotypic level, we obtained longitudinal samples from *NPM1*-mutant AML patients (n=8) undergoing 7+3 chemotherapy (**Extended Table 1, Extended Data Fig. 1**). Profiling of longitudinal samples revealed variable patterns in clonal evolution but most patients displayed notable alterations in the number of mutations and clones from diagnosis to complete response and/or relapse (**Fig. 4a; Extended Data Fig. 4a**). Interestingly, even in patients whose mutations remained the same, the distribution of mutant clones fluctuated throughout therapy. Our previous studies and others have identified significant genotype-immunophenotype correlations in *RAS* and/or *FLT3*-mutant clones^8,9,12^. In our cohort, we had two patients in particular who gained or lost signaling mutations while on therapy. In one patient, a dominant *NRAS*-mutant clone harboring co-occurring *TET2/NPM1* mutations was lost during therapy (Pt G; **Fig. 4bc; Extended Data Fig. 4b**). We found that the loss of this triple mutant clone in the relapse sample correlated with increased expression of dendritic and monocytic markers CD135 (FLT3) and CD16, respectively (CD135 *P* = 1.1 x 10^-209^, CD16 *P* = 9.6 x 10^-15^; **Fig. 4de**). The clonal evolution in Pt G was also correlated with decreased expression of myeloid and stem/progenitor markers CD33 and CD117 (CD33 *P* = 7.2 x 10^-211^, CD117 *P* = 5.2 x 10^-57^; **Fig. 4e; Extended Data Fig. 4cd**), Conversely, in a second patient, separate *RAS* and *FLT3*-mutant subclones were acquired upon relapse (Pt I; **Fig. 4fg; Extended Data Fig. 4e**). We found that compared to the diagnosis sample, the relapsed disease was enriched for higher expression of CD117 across the sample but also CD14 (CD117 *P* = 9.3 x 10^-31^, CD14 *P* = 5.0 x 10^-235^; **Fig. 4h; Extended Data Fig. 4f**). These findings confirm previously identified genotype-immunophenotype relationships and further suggest that 7+3 therapy can have varying effects on clonality and immunophenotype. Moreover, we have observed the profound alterations that gains/losses of signaling mutations have on the immunophenotype of a patient’s disease during therapy.

**Fig. 4.**
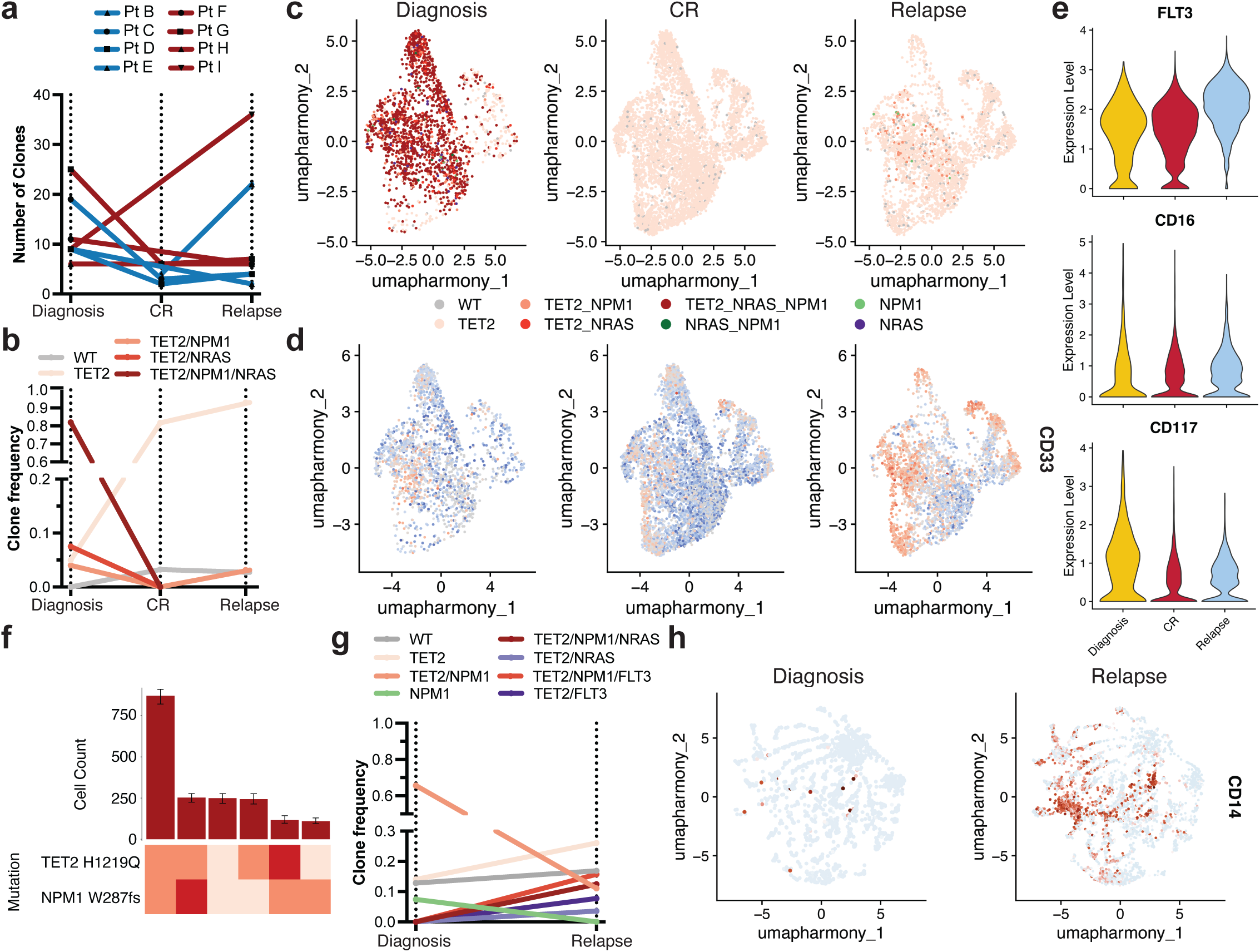
Clonal and immunophenotypic single cell analysis of longitudinal patient samples while undergoing 7+3 chemotherapy. **a)** Changes in number of clones for *NPM1*-mutant patients (n=8) where samples were analyzed longitudinally while undergoing therapy. Individual patients are indicated by connecting line with point at each disease state for which sample was available. Blue = *IDH2* co-mutation at diagnosis by bulk sequencing (n=4 patients); Red = *TET2* co-mutation at diagnosis by bulk sequencing (n=4 patients). **b-e)** Analysis of paired samples of representative patient G (Pt G) that underwent clonal change during treatment. **b**) Changes in clone frequencies at each disease state. Only genotypes identified in 1% or higher of total cells from at least one sample are depicted for clarity. Color of line denotes specific genotype also used in Fig. 4c. **c)** Uniform manifold approximation and projection (UMAP) plot of Pt G samples at diagnosis (left), CR (center), and relapse (right) clustered by immunophenotype with genotype overlaid. **d)** UMAPs of Pt G samples denoting relative expression of CD135 (FLT3) as patient underwent therapy. Color depicts relative expression (blue = low, red = high). **e)** Violin plots of selected immunophenotype markers (CD135/FLT3, top; CD16, center; CD117, bottom) that change significantly from diagnosis to relapse in Pt G samples. Color denotes disease state (diagnosis, yellow; CR, red; relapse, blue). **f-h)** Analysis of paired samples of representative patient I (Pt I) that underwent clonal change during treatment. **f)** Clonograph of diagnosis sample from patient I. Height of each bar represents the cell count of the clone identified below. Clone genotype is depicted by color with WT in light beige, heterozygous mutations in orange, and homozygous mutations in red. **g**) Changes in clone frequencies at each disease state as in Fig. 4b. **h**) Uniform manifold approximation and projection (UMAP) plots of Pt I samples at diagnosis (left) and relapse (right) clustered by immunophenotype with relative expression of CD14 overlaid. Color denotes relative expression (blue = low, red = high).

### Significant clonal evolution is correlated with genotypic, transcriptional, and immunophenotypic alterations

Our patient cohort also included a patient (Pt F) who displayed significant clonal evolution, i.e. a clonal sweep where the genotype and clones of the leukemia significantly changed^8^ (**Fig. 5a; Extended Data Fig. 5a**). This patient was initially diagnosed with AML that harbored *TET2 and NPM1* co-mutations. However, upon relapse on 7+3 therapy the patient instead harbored an *IDH1* mutation co-occurring with the *NPM1* mutation with no evidence of the initial *TET2*-mutant clones (**Fig. 5a, Extended Data Fig. 5a**). scProtein analysis revealed that while diagnosis clones were enriched for expression of monocytic markers CD14, CD11b, and CD33 (all *P* values *P* < 1.0 x 10^-250^), clones identified at relapse were enriched for CD117 expression suggesting a more immature phenotype following therapy (CD117 *P* = 8.24 x 10^-11^; **Fig. 5bc, Extended Data Fig. 5bc**). We next sought to connect these immunophenotypic changes to the clones identified at each disease state. We found genotype-immunophenotype relationships consistent with our cohort analysis (**Fig. 3d**). *TET2*-mutant clones at diagnosis showed increased expression of monocytic markers CD14, CD11b, CD16, and CD64 (adj *P* value for all < 1.0 x 10^-80^; **Fig. 5d, Extended Data Fig. 5d**). Conversely, *IDH1*-mutant clones at relapse showed decreased expression of monocytic markers in favor of CD117 (adj *P* value = 8.2 x 10^-11^). These findings reconfirm the existence of genotype-immunophenotype relationships and that they can drastically alter the immunophenotype of a patient’s disease based on the gain or loss of certain mutations and clones during therapy. Notably, these findings indicate that relapse can manifest in a more immature cell state, in contrast to numerous reports indicating monocytic differentiation is a therapy escape mechanism^22-24^.

**Fig. 5.**
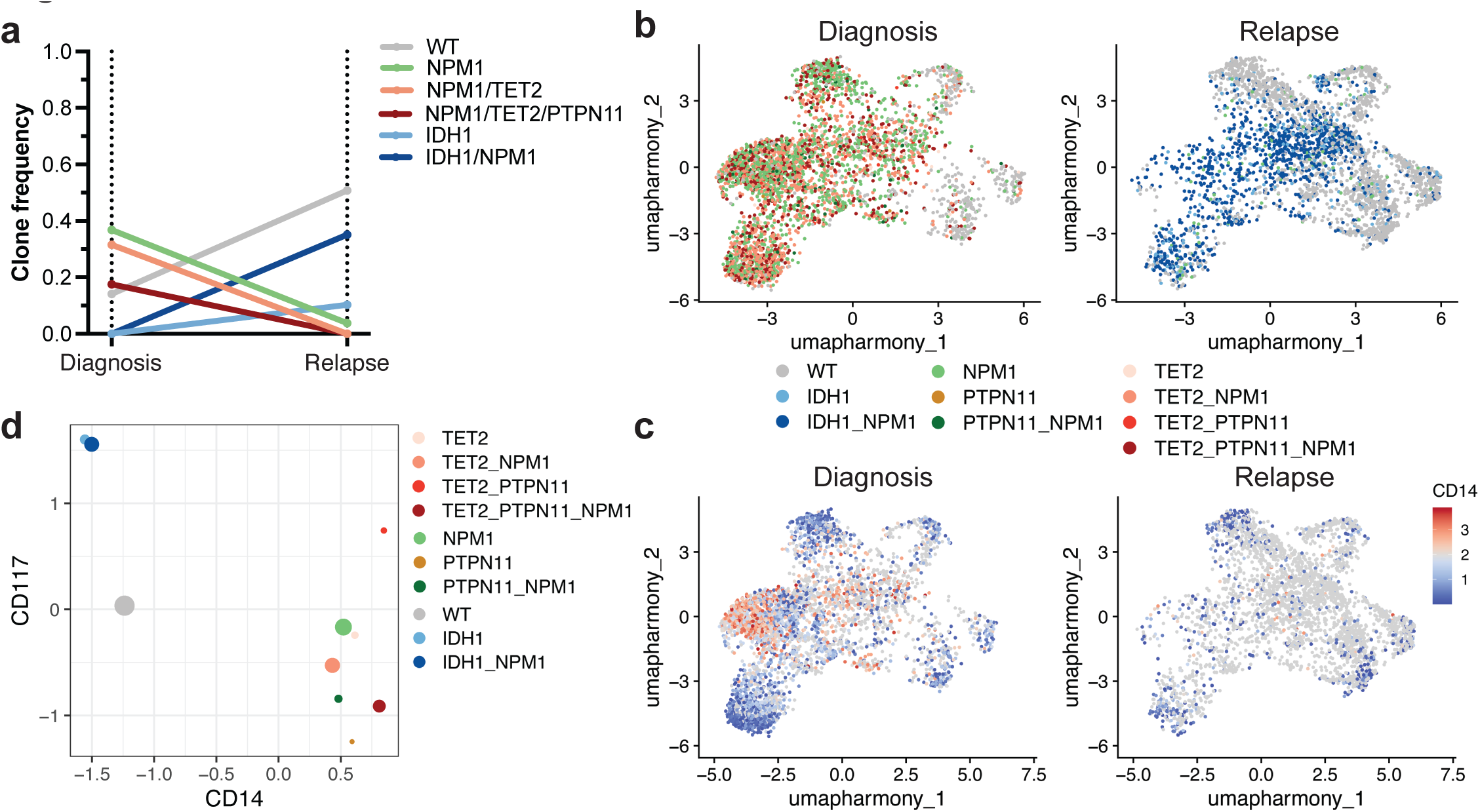
scDNA+Protein analysis of clonal sweep in Patient F. **a)** Changes in clone frequencies at each disease state. Only genotypes identified in 1% or higher of total cells from at least one sample are depicted for clarity. The color of line denotes specific genotype also used in Extended Data Fig. 5a. **b)** Uniform manifold approximation and projection (UMAP) plot of Pt F samples at diagnosis (left) and relapse (right) clustered by immunophenotype with genotype overlaid. **c)** UMAP from **b** with relative expression of CD14 overlaid. Color depicts relative expression (blue = low, red = high). **d)** Plot depicting expression of immunophenotypic markers CD117 (Y axis) and CD14 (X axis) by each identified clone found in Pt F samples. Normalized expression of each marker is depicted by dot location with size of dot denoting the fraction of the clone that expresses the marker. Genotype is denoted by same color as in Fig. 5a and 5b.

To connect these immunophenotype changes to gene expression changes, we performed single cell RNA-seq with cellular indexing of transcriptomes and epitopes (CITE-seq) analysis of the AML samples. We queried the matched samples from Pt F (**Fig. 5**), which showed drastic clonal evolution and an *IDH2*-mutant relapse sample (Pt B) whose *IDH2* mutation was stable from diagnosis and relapse (**Fig. 6**). First, we identified captured cell types through label transfer from a normal adult hematopoiesis reference^25^. These AML samples contained cell clusters identified as Multilineage/GMPs (Multilin-GMP-1), monocytes (Intermediate Mono, Classical/Non-classical Mono, Mono), dendritic cells (pre-DC, cDC), and T cells (CD8, CD4) (**Fig. 6a; Extended Data Fig. 6a**). Compared to the diagnosis sample, we identified a decrease in Intermediate Mono-1 and Intermediate Mono-2 cells and an increase in Multilin-GMP-1 cells in relapsed samples (**Extended Data Fig. 6b**). We subsequently focused on immunophenotype changes within the three samples across the three most prevalent cell clusters (**Extended Data Fig. 6c**). A subset of Multilin-GMP cells in the relapse samples trended towards increased CD117 expression compared to the diagnosis sample (Pt F relapse: did not reach significance; Pt B relapse: P<0.0002; **Fig. 6b**), aligning with our findings of *IDH2*-mutant clones and heightened expression of stem/progenitor markers. Moreover, a prominent intermediate monocyte population (Intermediate Mono 1) showed increased expression of CD14 and CD11b in the Pt F diagnosis sample compared to the paired relapse sample (CD14 *P* < 0.0001; CD11b *P* = 0.0008; **Fig. 6cd**). When we evaluated the gene expression of shared cell surface markers between our CITE-seq and scDNA+Protein immunophenotype panels, we found that expression of certain marker genes like *CD14* and *ITGAM* (CD11b) correlated well with protein expression, whereas other genes like *IL3RA* (CD123) had stable RNA expression compared to variable cell surface expression across the three clusters (**Extended Data Fig. 6cd**). Overall, these findings suggest that CITE-seq can refine the immunophenotype patterns observed in our scDNA+Protein analysis to specific cell populations.

**Fig. 6.**
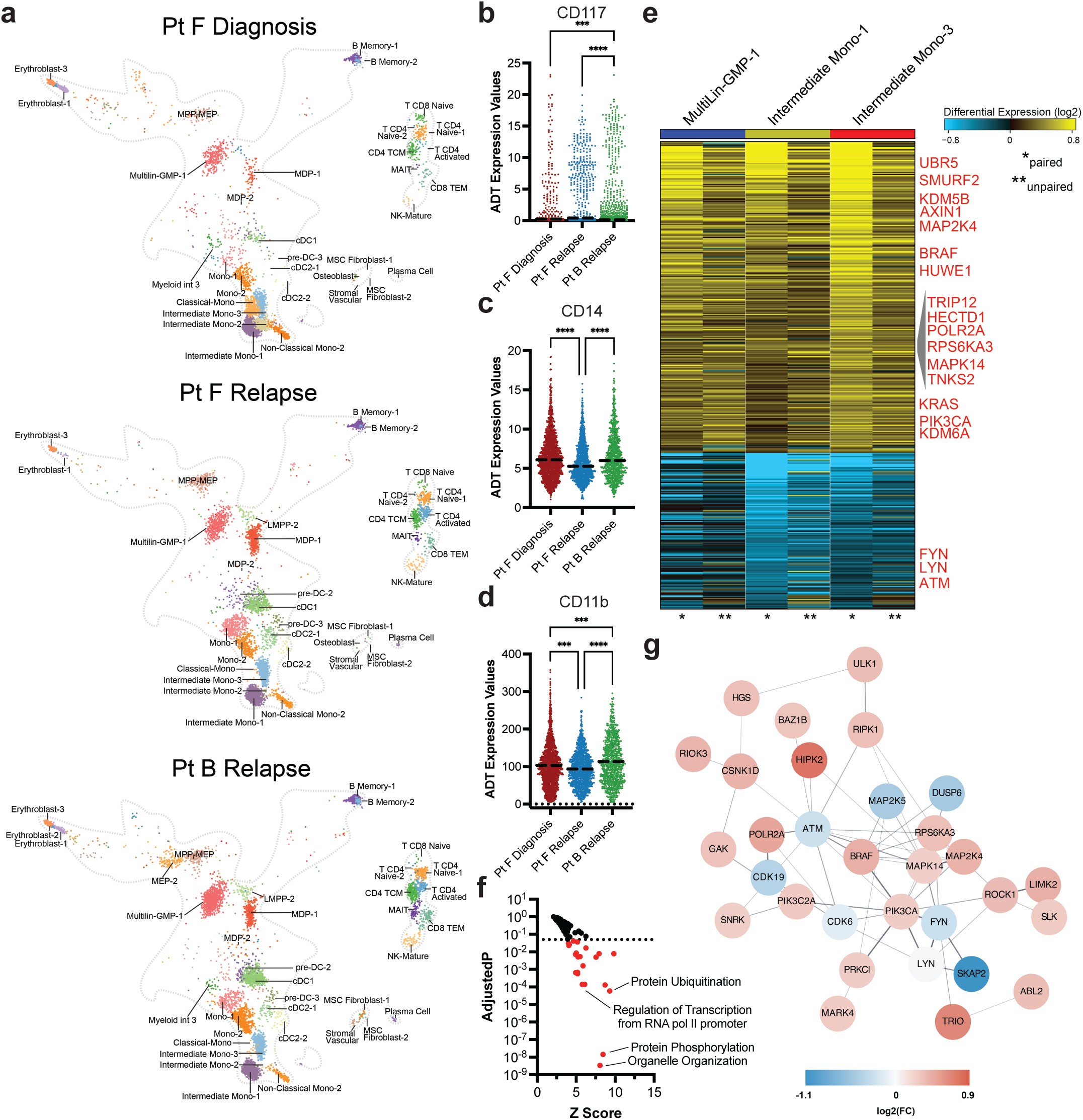
Matched CITE-seq analysis correlates scDNA+Protein results and identifies differentially expressed pathways upon relapse. **a)** UMAPs derived from CITE-seq analysis and clustered based on similarities to reference cell clusters (Extended Data Fig. 6a) for Pt F diagnosis (top), Pt F relapse (center), and Pt B relapse (bottom). Cell cluster identities are denoted based on cell identity in reference atlas (Extended Data Fig. 6a). **bcd)** Scatter dot plots of single cell CD117 (**b**), CD14 (**c**) and CD11b (**d**) antibody tag reads from cells clustered as Multilin-GMP-1 (**b**) or Intermediate Mono-1 (**cd**) cells from each sample (n=3). Bold dotted line denotes the mean. Kruskal-Wallis test was used to determine statistical significance amongst groups. ***P<0.001, ****P<0.0001 denoted. **e**) Heatmap of genes found to be differentially expressed in Multilin-GMP-1 (left, blue bar), Intermediate Mono-1 (center, yellow bar), and Intermediate Mono-3 (right, red bar) cell clusters between Pt F Diagnosis and both relapse samples (Pt F Relapse, Pt B Relapse). Heatmap scale denotes log fold gene expression from high (yellow) to low (blue). Select genes are denoted in red. Asterisks indicate whether column is a comparison between the paired samples (* denotes Pt F Diagnosis-Pt F Relapse) or unpaired samples (** denotes Pt F Diagnosis-Pt B Relapse). **f)** Waterfall plot of Z-scores (X) and adjusted P values (Y) for GO:Biological Processes found to be differentially expressed in Pt F diagnosis sample compared to Pt F and Pt B relapse samples by AltAnalyze. Color of dot denotes significance based on adjusted P values < 0.05 (dashed line) in red. **g)** Network connectivity map denoting the interactions of differentially expressed genes from Fig. 6e.

We next interrogated significant gene expression changes between diagnosis and relapse, focusing on the cell populations which correlated with our scProtein immunophenotypic alterations. Upon comparing our diagnosis sample to the patient paired (Pt F) and unpaired (Pt B) relapse samples, we found significant differential expression of several protein ubiquitination genes including upregulation of *HUWE1* and *HECTD1*, E3 ubiquitin ligases with established roles in leukemia^26^ and stem cell^27^ proliferation and regeneration, respectively (**Fig. 6ef**). Multiple genes involved in Wnt/β-catenin pathway activation, previously shown to be important in *MLL*-rearranged and *HOX*-dependent leukemia development^28-30^, were upregulated in relapse samples, including *AXIN1*, *LRRFIP2*, and *UBE2B*. Protein interaction analysis uncovered a significant upregulation of multiple kinase and phosphatase genes including *KRAS*, *BRAF,* and *PIK3CA,* suggesting transcriptional changes to the RAS-MAPK-PI3K pathways and other signaling networks, known to play roles in development of therapy resistance^24,31,32^ (**Fig. 6fg**). Further, *SMURF2* was found to be upregulated as well in relapse samples, which has been previously implicated in controlling KRAS protein stability^33^. RAS/MAPK pathway activating mutations have been commonly found at relapse from various targeted therapies^12,23,34,35^. Our results indicate that there is substantial gene expression dysregulation of signaling cascades, including RAS/MAPK, even in the absence of activating mutations. Collectively, these studies indicate that the genotypes of AML clones play significant roles in dictating the cellular immunophenotypes and clonal lineage potentials, underscoring the need for further resolution of genotype-transcriptome-immunophenotype relationships in AML development and evolution.

## Discussion

*NPM1* is one of the most commonly mutated genes in AML^1-3,6^, and previous studies have suggested different levels of synergy between *NPM1* and co-occurring mutations^8,9,36^, as well as significant clonal changes while patients are undergoing targeted therapies^12,13^. In this study, we have utilized scDNA+Protein analysis to further examine the clonal architecture patterns in *NPM1-* mutated AML samples, as well as understand genotype-immunophenotype correlations as patients undergo standard of care chemotherapy. We first identified that differences in clonal architecture exist depending on the co-occurring mutations with *NPM1* on a patient level. Notably, we also found that *RAS/FLT3*-mutant AMLs had significantly increased clonal diversity, particularly in the relapse setting, suggesting that AMLs with signaling gene mutations may use clonal heterogeneity to drive relapse compared to AMLs without signaling genes. The findings in our study could be critical in understanding why further insight into these mutations and mutational combinations holds importance for even more nuanced risk stratification for AML patients. While our studies here focus on a very common subtype of AML with co-mutations that exist broadly across all AML patients, the scDNA targeted amplicon panel does exclude the identification of all possible mutations that exist and/or are gained and lost during therapy. The mutational cooperativity analyses in our study are important in helping to understand differences in clone sizes; however, our cooperativity results may be limited by potential allele dropout in *NPM1* single-mutant clones. Furthermore, additional non-somatic alterations may be playing a role in leukemogenesis and clonal evolution^37,38^, which could be further explored by scRNA sequencing and scATAC-seq.

Additionally, our single cell immunophenotypic analyses in this study revealed specific genotype-immunophenotype relationships. Surprisingly our data suggests that mutations thought of as initiating mutations, (*DNMT3A*, *TET2*, and *IDH1/2*), seem to dictate the lineage trajectories for subsequent clones. For instance, we found that *DNMT3A*- and *IDH*-mutant clones, with or without *NPM1* and signaling mutations, show enrichment in hematopoietic stem/progenitor cell markers CD117, CD123, and CD34. Meanwhile, *TET2*-mutant clones show increased expression of monocytic markers CD14, CD11b, and CD64. The remarkable divergence between *DNMT3A* and *TET2*, the two most frequent mutations found in clonal hematopoiesis^39^, suggests that the immunophenotypes of the mutant leukemic clones may be influenced even by these early mutations during leukemogenesis. Moreover, our studies infer that while initiating mutations provide possible lineage trajectories, signaling mutations can refine these trajectories, again underscoring that the genotype-immunophenotype relationships are highly unique to the combination of mutations. Further studies are needed to understand the importance of these relationships and how they impact response to both cytotoxic and targeted therapies.

Our study found notable clonal and immunophenotypic changes from diagnosis to relapse as patients underwent standard cytotoxic chemotherapy. Relapse and refractory disease are major contributors to the dismal outcomes observed in AML patients, with a 5-10% 5-year survival rate in patients with relapsed/refractory disease^40^. A better understanding of resistance mechanisms and leukemic evolution as patients undergo therapy, can influence clinical management and therapeutic options for AML patients. Interrogating longitudinal samples from patients who underwent 7+3 therapy, we found that most patients’ disease expressed more of an immature phenotype in relapse, previously suggested in small studies of AML samples^41^. This is in contrast with the changes that are observed the combination of the BCL-2 inhibitor, venetoclax, and hypomethylating agent, azacitidine (Ven/Aza) and other recent therapies. A recurrent mechanism of acquired resistance/relapse for Ven/Aza lies in the expansion of a myelomonocytic phenotype blast population, characterized by higher CD11b/CD14 expression and enriched for *NRAS/MAPK* mutations^22-24,34^. While outside the scope of this study, these divergent findings bring into question whether the selective pressures imposed by different treatment regimens influence how leukemic clones respond and therefore how clonotypes and immunophenotypes will change during therapy and upon relapse.

Lastly, we performed CITE-seq on matched diagnosis and relapse samples from two patients in our cohort, one of whom displayed significant clonal evolution while on 7+3 therapy. In doing so, we could identify cell populations displaying the immunophenotype alterations correlating with the clonal evolution and uncover significant gene expression changes. These gene expression changes suggest that dysregulation of signaling pathways and ubiquitination pathways can play a role in clonal evolution while on therapy. Not surprisingly, *RAS*-*MAPK-PI3K* pathways were among the significantly upregulated pathways, which align with many recent studies of resistant disease^12,23,34,35^. Notably, neither of the relapse samples harbored or acquired *RAS/MAPK* signaling mutations but instead upregulated the intrinsic pathway through transcriptional alterations. While this analysis was limited to a small number of samples, these findings highlight the need to understand how clones are evolving both at the genotype and immunophenotype level, but also at the transcriptomic level. Truly integrated trimodal analysis of genotype, transcriptome, and immunophenotype is yet to be obtained, but will likely provide a new level of understanding of how mutations synergize to drive leukemic development and disease progression.

Analyzing clonal evolution at a single-cell level provides insights into how *NPM1* mutations cooperate with epigenetic and signaling mutations to generate clonal complexity, underlying resistance to treatment. Matched CITE-seq analysis suggests widespread changes to biological processes including signaling and protein ubiquitination pathways. These studies nominate dynamic pathway changes that might contribute to disease relapse. Collectively, our investigation underscores the need to further study AML patients longitudinally and at high cellular resolution, to discover mechanisms of response and relapse to current therapies. We anticipate that similar integrated multiomic approaches will enable new risk stratifications that predict treatment responses and inform therapeutic strategies that target cancer as an evolving, multi-clonal disease.

**Extended Table 1.** Single cell sequencing analytical metrics for each sample in the *NPM1*-mutant AML cohort (n=43). Values provided by the Mission Bio Tapestri pipeline after initial processing. Longitudinal samples from the same patient are denoted with sample name with an underline and disease state annotation (diagnosis= d, complete response= cr, relapse= r). PBMC = peripheral blood mononuclear cells; BM = bone marrow.

| Sample Name | Cell count | Panel Uniformity | Mean read/cell/amplicon | %DNA read pairs assigned to cell | Mean reads/cell/antibody | Sample Type |
| --- | --- | --- | --- | --- | --- | --- |
| A | 11,402 | 93% | 67 | 76% | 193 | PBMC |
| AA | 5,082 | 79% | 91 | 22% | 297 | BM |
| AB | 40,663 | 93% | 58 | 75% | 160 | PBMC |
| AC | 9,939 | 80% | 13 | 13% | 108 | BM |
| AD | 9,429 | 93% | 95 | 63% | 166 | BM |
| AE | 31,027 | 94% | 55 | 75% | 73 | BM |
| AF | 45,758 | 94% | 72 | 78% | 147 | PBMC |
| B_cr | 7,631 | 94% | 87 | 78% | 312 | PBMC |
| B_r | 13,936 | 95% | 38 | 69% | 177 | BM |
| C_cr | 12,865 | 95% | 44 | 78% | 321 | BM |
| C_d | 9,669 | 94% | 55 | 72% | 468 | BM |
| C_r | 4,248 | 94% | 207 | 66% | 421 | BM |
| D_cr | 10,218 | 96% | 35 | 58% | 195 | PBMC |
| D_d | 7,112 | 92% | 62 | 48% | 119 | PBMC |
| D_r | 8,726 | 94% | 60 | 51% | 175 | PBMC |
| E_d | 8,442 | 92% | 69 | 52% | 189 | PBMC |
| E_r | 1,184 | 90% | 541 | 54% | 491 | PBMC |
| F_d | 8,175 | 94% | 205 | 74% | 611 | BM |
| F_r | 7,036 | 94% | 100 | 71% | 344 | BM |
| G_cr | 15,545 | 95% | 55 | 76% | 162 | BM |
| G_d | 6,414 | 95% | 178 | 77% | 379 | BM |
| G_r | 9,427 | 95% | 69 | 74% | 250 | BM |
| H_d | 7,522 | 94% | 78 | 51% | 188 | BM |
| H_r | 10,404 | 92% | 64 | 52% | 98 | BM |
| I_d | 2,562 | 93% | 255 | 55% | 546 | PBMC |
| I_r | 10,911 | 92% | 67 | 65% | 151 | PBMC |
| J | 9,556 | 92% | 82 | 57% | 139 | PBMC |
| K | 3,800 | 92% | 200 | 63% | 248 | PBMC |
| **L** | 17,317 | 94% | 75 | 68% | 187 | PBMC |
| **M** | 881 | 81% | 301 | 48% | 230 | PBMC |
| **N** | 2,051 | 90% | 283 | 58% | 483 | PBMC |
| **O** | 22,651 | 92% | 72 | 66% | 79 | BM |
| **P** | 17,451 | 93% | 109 | 67% | 50 | BM |
| **Q** | 27,726 | 93% | 74 | 65% | 73 | BM |
| **R** | 26,340 | 93% | 68 | 73% | 167 | BM |
| **S** | 31,160 | 93% | 39 | 61% | 183 | BM |
| **T** | 25,512 | 93% | 74 | 69% | 184 | PBMC |
| **U** | 6,315 | 94% | 102 | 77% | 468 | BM |
| **V** | 10,602 | 93% | 18 | 64% | 177 | BM |
| **W** | 35,124 | 94% | 74 | 69% | 184 | PBMC |
| **X** | 13,452 | 93% | 99 | 80% | 420 | BM |
| **Y** | 26,905 | 93% | 61 | 70% | 242 | BM |
| **Z** | 17,144 | 93% | 53 | 68% | 275 | BM |
| **Average** | **14,170** | **92%** | **104.74** | **64%** | **244.88** |  |
| **Total** | **609,314** |  |  |  |  |  |

**Extended Table 2.** Clinical characteristics and treatment information of patients with longitudinal samples analyzed by scDNA+Protein. Ara-C and Daunorubicin treatment is also known as 7+3 therapy.

| Patient | Induction Regimen | Age (y) | Sex | Race | Days from treatment start to CR | Days from treatment start to Relapse | Days from CR to Relapse |
| --- | --- | --- | --- | --- | --- | --- | --- |
| B | Ara-C, Daunorubicin, Etoposide | 54 | Male | White | 27 | 3044 | 3017 |
| C | Ara-C, Daunorubicin, Etoposide | 75 | Male | White | 39 | 415 | 376 |
| D | Ara-C, Daunorubicin, G3139 | 74 | Female | White | 41 | 606 | 565 |
| E | Ara-C, Daunorubicin | 60 | Male | White | 31 | 1484 | 1453 |
| F | Ara-C, Daunorubicin, Etoposide | 53 | Male | White | 35 | 914 | 879 |
| G | Ara-C, Daunorubicin, Etoposide | 29 | Female | White | 28 | 1099 | 1071 |
| H | Ara-C, Daunorubicin, Etoposide | 74 | Male | White | 39 | 136 | 97 |
| I | Ara-C, Daunorubicin, Etoposide | 60 | Male | White | 27 | 2175 | 2148 |

**Extended Data Fig. 1.**
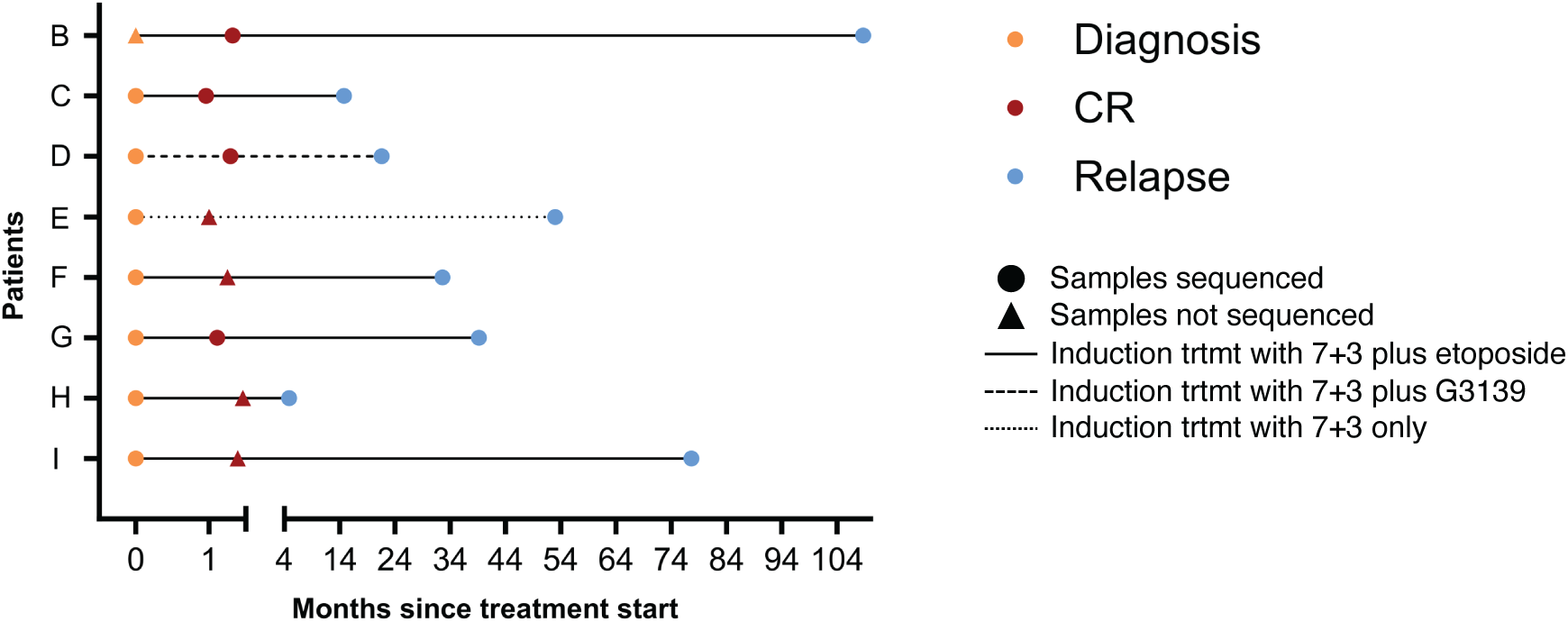
Treatment response courses for patients (n=8) while on 7+3 therapy. Each patient is labeled on Y axis with months since treatment start denoted on X axis. Diagnosis (yellow), complete response (red), and relapse (blue) samples that were available for sequencing denoted by colored circles with timepoints with unavailable samples depicted by triangles at time point based on location of dot. Therapy is denoted by line style (complete = 7+3 plus etoposide; large dash = 7+3 plus G3139; small dash = 7+3 alone). Patient outcomes are not provided or denoted on graph.

**Extended Data Fig. 2.**
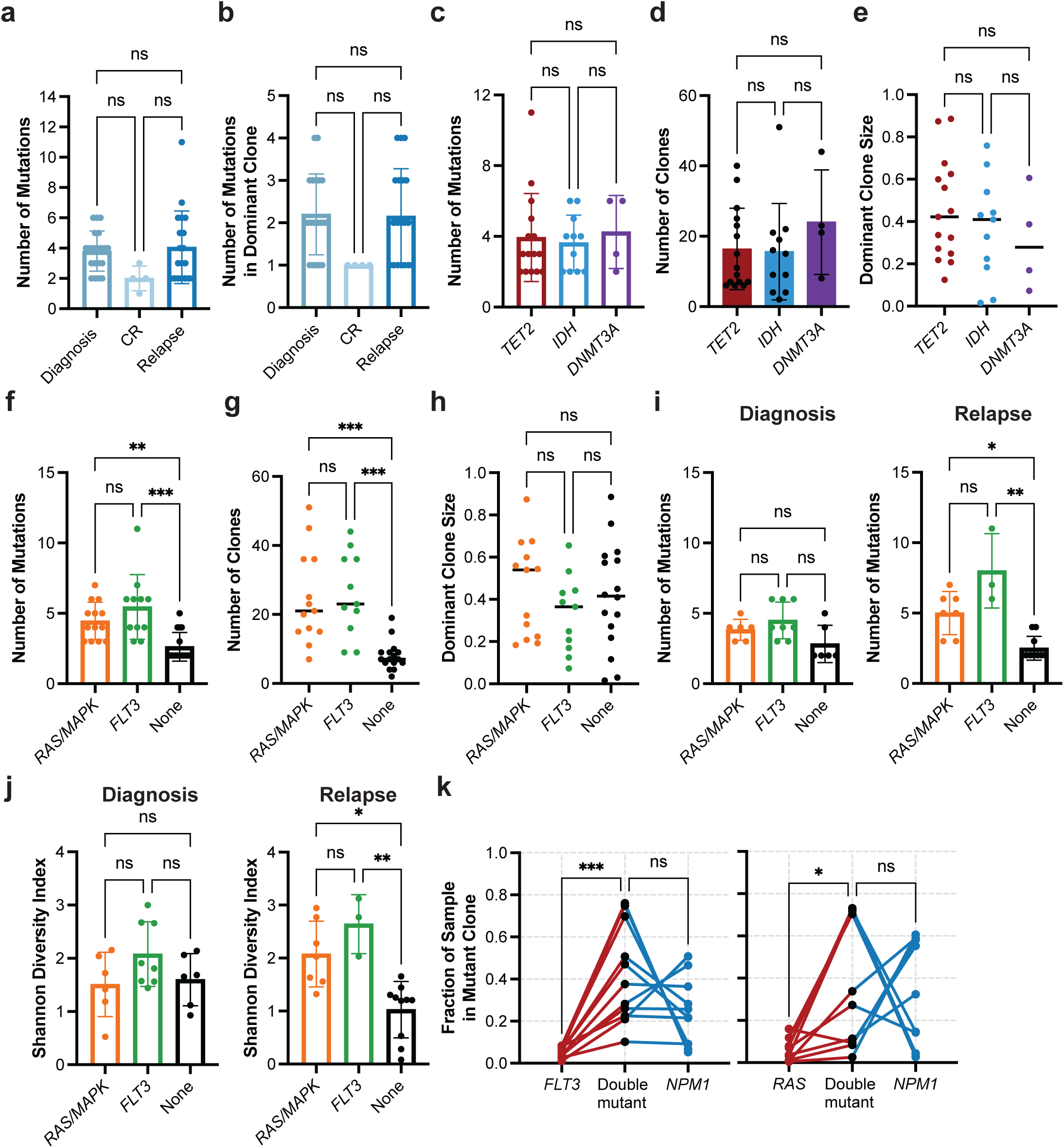
Analysis of clonal architecture by disease state and by gene mutation. **a-b)** Clonal architecture metrics for entire cohort (n=43 samples) by disease state, including (**a**) number of mutations per sample, and (**b**) number of mutations in the dominant clone. **c-e)** Bar graphs depicting clonal architecture metrics of samples (n=30) with different epigenetic gene mutations in *TET2* (red), *IDH1/2* (blue), or *DNMT3A* (purple) at diagnosis and relapse states, including (**c**) number of mutations per sample, (**d**) number of clones per sample, and (**e**) dominant clone size. **f-h)** Number of mutations per sample (**f**), number of clones per sample (**g**) and dominant clone size (**h**) for samples with *RAS/MAPK* (n=13; orange) or *FLT3* (n=11; green) mutations vs. no signaling gene mutations (n=16; None, black), at diagnosis and relapse states combined. **i**) Number of mutations per sample (as in Extended Data Fig 2f) stratified by diagnosis (left panel) or relapse (right panel). **j**) Clonal diversity, as calculated by Shannon diversity index, for samples with *RAS/MAPK* (n=13) or *FLT3* (n=11) mutations vs. no signaling gene mutations (n=16) at diagnosis and at relapse. **j)** Fraction of sample in single- and double-mutant clones in *FLT3-NPM1* (n = 11; left panel) and *RAS-NPM1* (n = 10; right panel) mutant samples. Individual samples denoted by connecting lines. **a-d, f, i-j)** Mean value for each cohort shown by height of bar with standard deviation depicted with error bars. **e, g-h)** Center line - median value for each cohort. Kruskal-Wallis test was used to determine statistical significance amongst groups for all panels except **k** where a two-way ANOVA was used. *P<0.05, **P<0.01, ***P<0.001 denoted for all panels.

**Extended Data Fig. 3.**
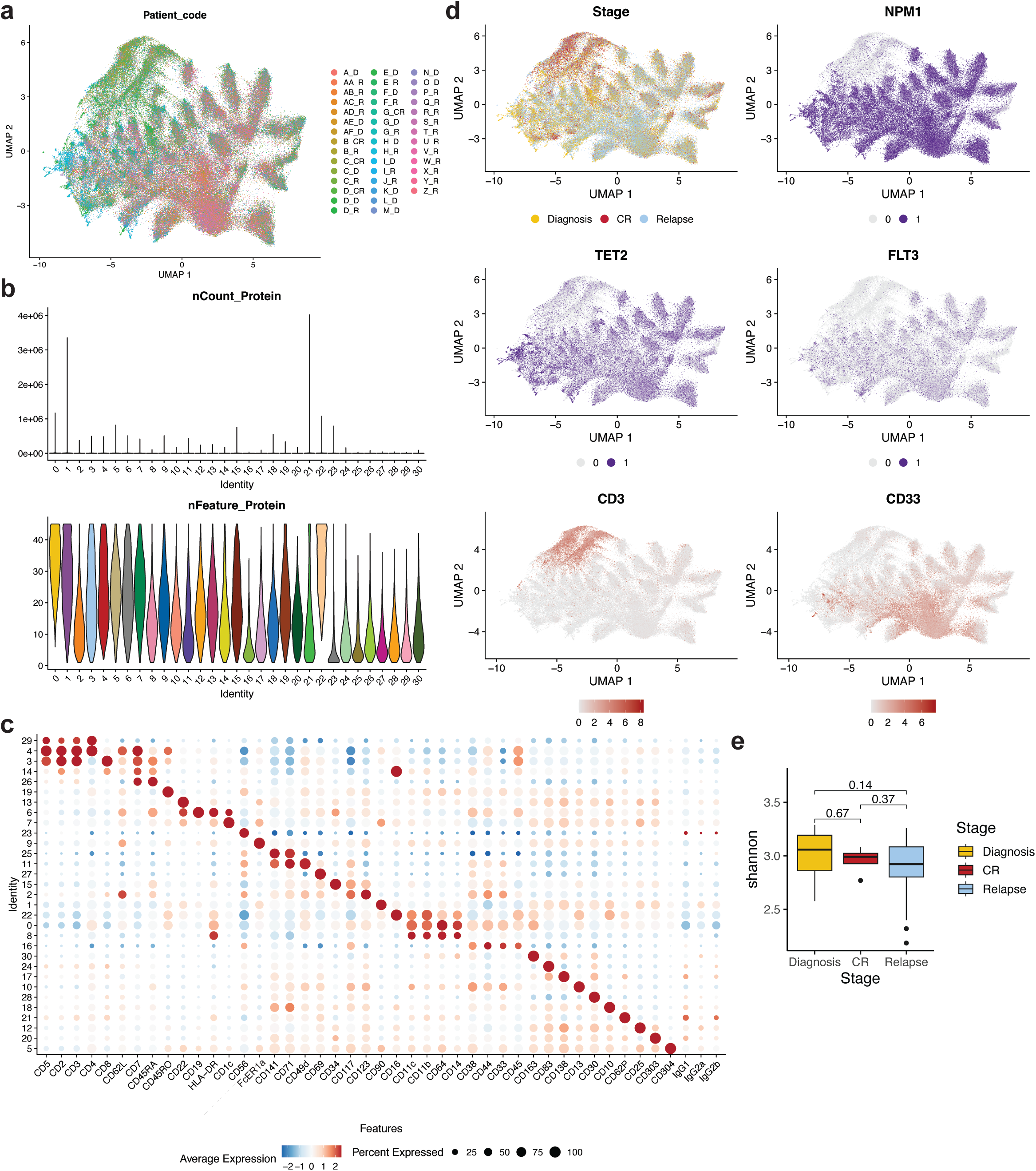
Immunophenotype analysis of all single-cell DNA+Protein samples. **a)** Uniform manifold approximation and projection (UMAP) plot of all patient samples (n=43) clustered by immunophenotype. Cells from the same patient sample are shown in the same color. **b)** Single cell immunophenotype metrics for each community denoted by total number of sequencing reads for each community (nCount Protein; top panel) and violin plot denoting number of unique proteins expressed in each community (nFeature Protein; bottom panel. Colors of each community in bottom panel match colors from Extended Data Fig. 3b. **c)** Dot plot depicting relative expression of each immunophenotypic marker within each community. Normalized expression of each marker depicted by color (blue = low, red = high) with size of dot denoting the fraction of cells within each community that expresses the marker. **d)** Uniform manifold approximation and projection (UMAP) plots of entire patient cohort (n=43) with cells clustered by immunophenotype. Top left, disease stage overlaid onto the UMAP (Diagnosis, yellow; Complete response, CR, Red; Relapse, blue). Top right panel (NPM1) and middle panels (TET2, right; FLT3, left), select mutant genes overlaid onto the UMAP (wildtype, grey; mutant, purple). Bottom panel, select immunophenotypic markers (CD3, left panel; CD33, right panel) overlaid onto the UMAP with expression from low (grey) to high (red) expression depicted. **e)** Box and whisker plot of community diversity within each disease stage (diagnosis, yellow; CR, red; relapse, blue) calculated by Shannon diversity index.

**Extended Data Fig. 4.**
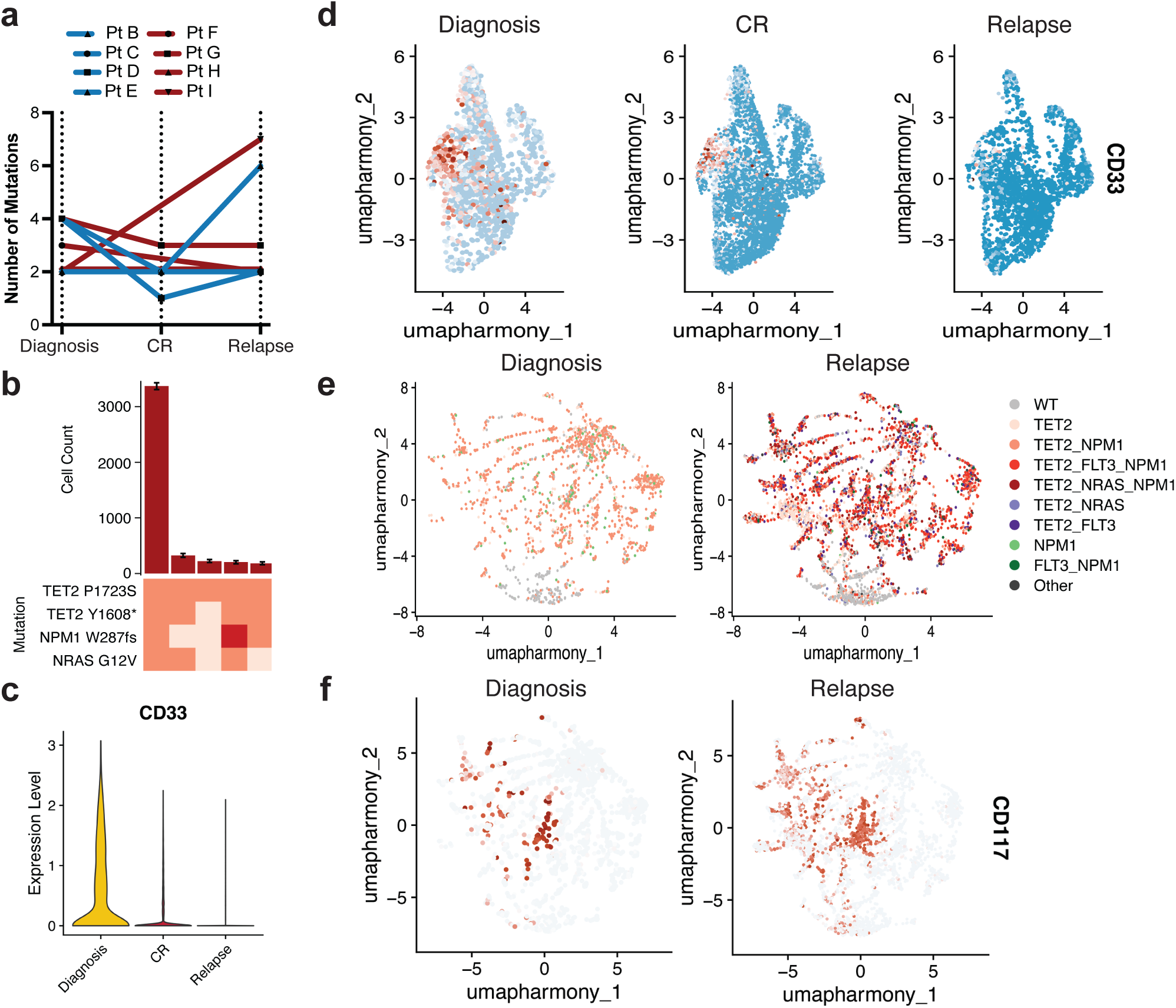
Alterations in clonality and immunophenotype during 7+3 therapy. **a)** Changes in number of mutations for *NPM1*-mutant patients (n=8) where samples were analyzed longitudinally while undergoing therapy. Individual patients indicated by connecting line with point at each disease state for which sample available. Blue = *IDH2* co-mutation at diagnosis by bulk sequencing (n=4 patients); Red = *TET2* co-mutation at diagnosis by bulk sequencing (n=4 patients). **b)** Clonograph of diagnosis sample from Pt G. Height of each bar represents the cell count of the corresponding identified clone noted below. Clone genotype is depicted by color with WT (light beige), heterozygous (orange), and homozygous (red) mutations denoted. **c)** Violin plot of CD33 in Pt G samples. Color denotes disease state (diagnosis, yellow; CR, red; relapse, blue). Bold dotted line denotes the mean with quartiles shown by thin dotted lines. **d)** UMAPs of Pt G samples denoting relative expression of CD33 as patient underwent therapy. Color depicts relative expression (blow = low, red = high). **e-f)** UMAP plots of Pt G samples at diagnosis (left), CR (center), and relapse (right) clustered by immunophenotype with genotype (**e**) or relative expression of CD117 (**f**) overlaid. Colors in **e** denote genotype colors in Fig. 4g. Colors in **f** denote relative expression of CD117 (blue = low, red = high).

**Extended Data Fig. 5.**
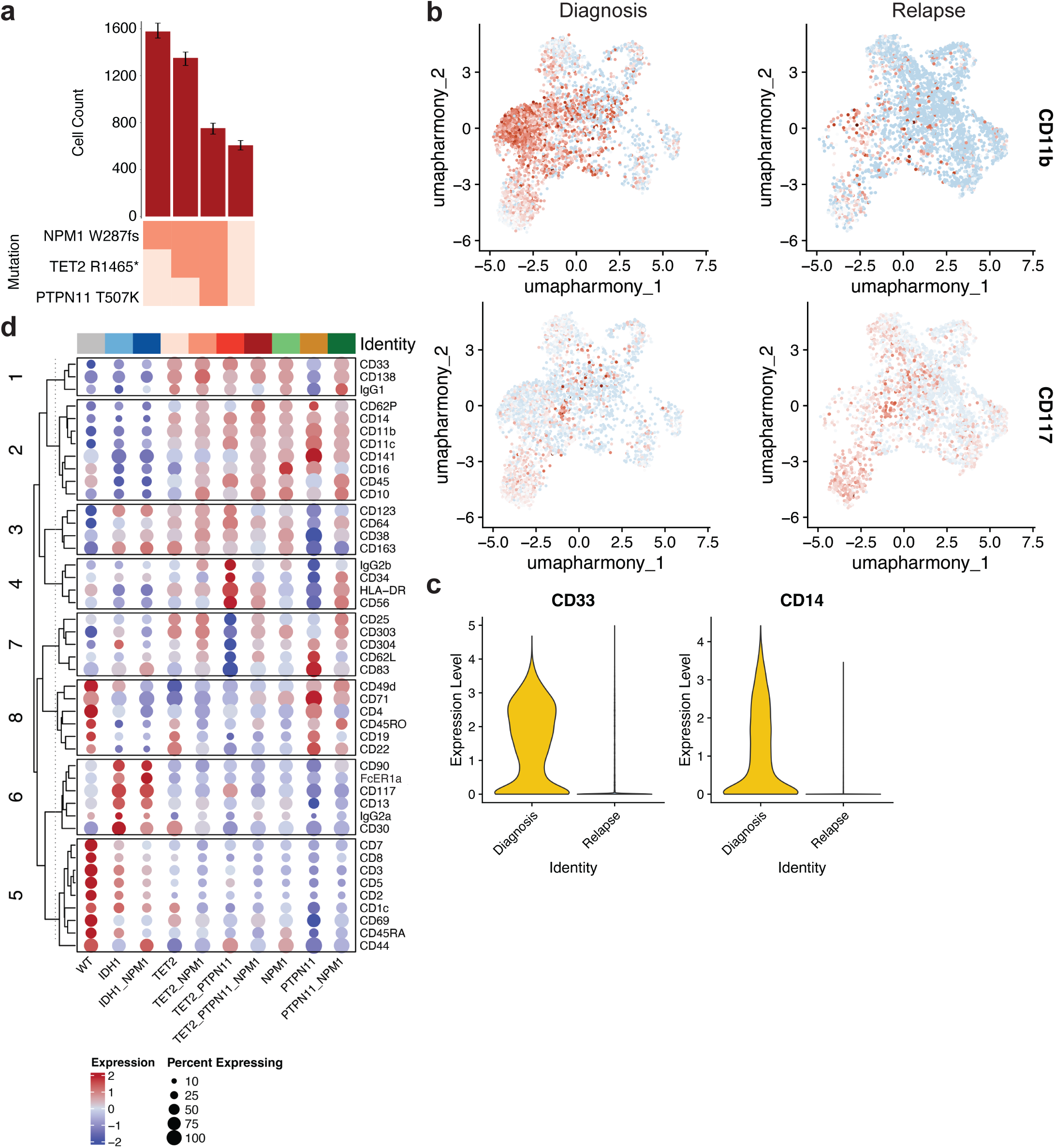
Clonal sweep during 7+3 therapy. **a)** Clonograph of diagnosis sample from patient F. Height of each bar represents the cell count of the corresponding identified clone noted below. Clone genotype is depicted by color with WT (light beige), heterozygous (orange), and homozygous (red) mutations denoted. **b)** UMAPs from Fig. 5b with relative expression of CD11b (top) and CD117 (bottom) overlaid. Color depicts relative expression (blue = low, red = high). **c)** Violin plot of CD33 (right panel) and CD14 in Pt F samples. Color denotes disease state (diagnosis, yellow; relapse, blue). Bold dotted line denotes the mean with quartiles shown by thin dotted lines. **d)** Dot plot depicting expression of immunophenotypic markers by genotype-specific clones identified in Pt F samples. Normalized expression of each marker depicted by color (blue = low, red = high) with size of dot denoting the fraction of cells within each genotyped clone that expresses the marker. Immunophenotype markers grouped by corresponding lineage associations. Full genotype for each row denoted at left of the dotplot.

**Extended Data Fig. 6.**
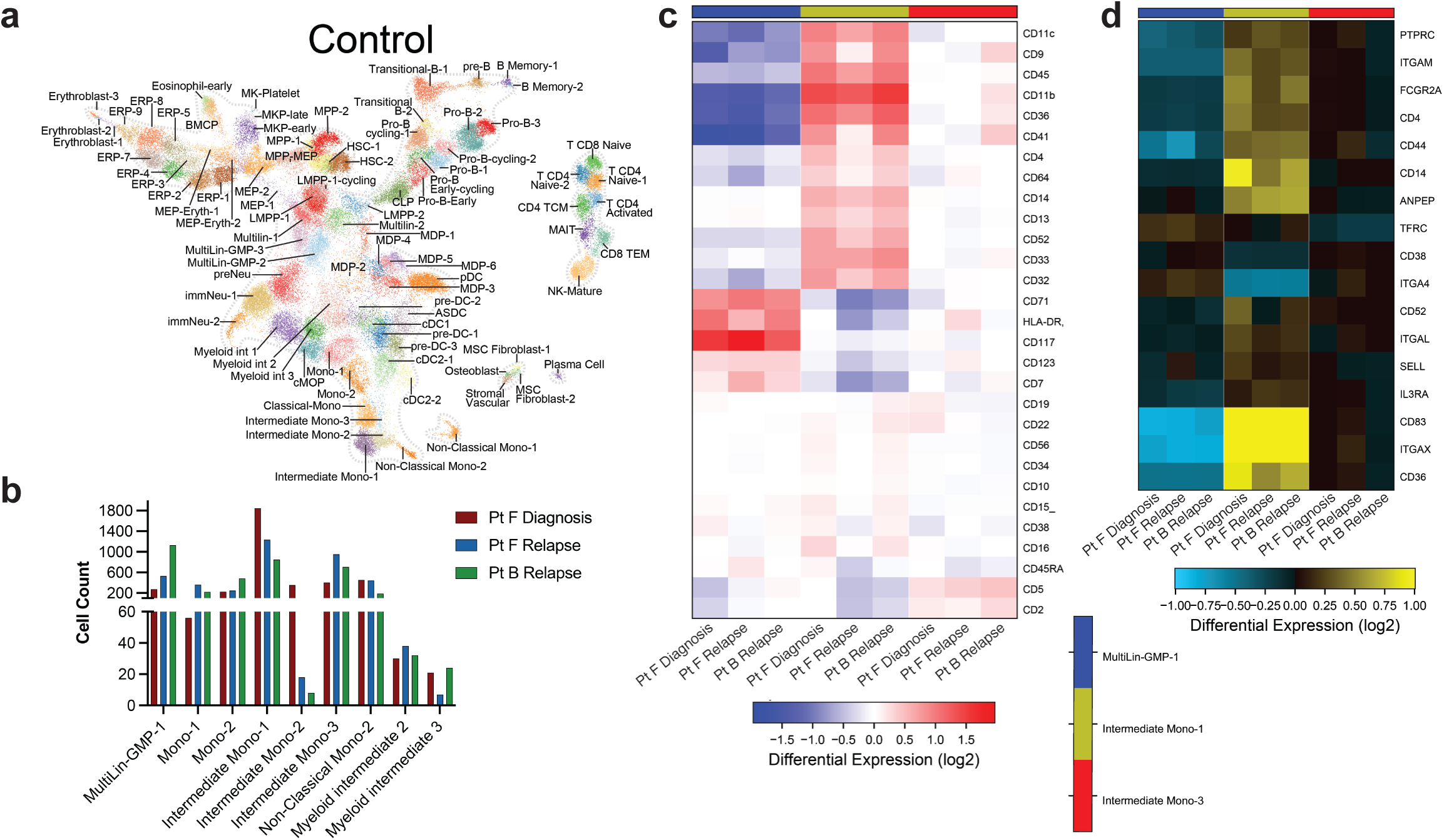
CITE-seq analysis of clonal evolution. **a)** UMAP cell cluster atlas^25^ of human hematopoiesis derived from CITE-seq analysis and used as reference map for Pt F and Pt B samples in Fig. 6. Cell cluster identities denoted. **b)** Bar plot of cell counts for selected cell clusters identified from CITE-seq analysis for samples analyzed (n=3). Color of bar denotes the sample identity with legend. **c)** Heatmap of cell surface marker ADT read counts for antibodies used in the CITE-seq panel across Multilin-GMP-1 (left, blue column), Intermediate Mono-1 (center, yellow column), and Intermediate Mono-3 (right, red column) cell clusters. Heatmap scale denotes log fold differences in read counts from high (red to low (blue). **d)** Heatmap of cell surface marker gene expression for antibodies used in either CITE-seq and/or scDNA+Protein panel across Multilin-GMP-1 (left, blue column), Intermediate Mono-1 (center, yellow column), and Intermediate Mono-3 (right, red column) cell clusters. Only differentially expressed genes are included. Heatmap scale denotes log fold gene expression from high (yellow) to low (blue).

## Methods

### Reagents

Tapestri related reagents were included as part of the Myeloid Clonal Evolution DNA+Protein sequencing kit purchased from Mission Bio with the following exceptions: TotalSeqD Antibody Cocktail v2, Cell Staining Buffer, TotalSeqD CD135 antibody were purchased from Biolegend. The Myeloid Clonal Evolution amplicon panel has been described previously^8^.

### Patient samples

Patient consent was obtained according to protocols approved by the Institutional Review Boards in accordance with the Declaration of Helsinki. This study was approved by CCHMC IRB (protocol 2022-0806), MSKCC IRB (protocol #15-017), and OSU IRB (#2023C0062). WHO classification criteria were used for diagnosis and disease status assignment^4^. Patient samples were collected and processed by institutional biorepositories. Peripheral blood or whole bone marrow mononuclear cells were isolated by centrifugation on Ficoll and viably frozen. High-throughput genetic sequencing was utilized to profile each sample. MSKCC samples were profiled using HemePACT, a targeted deep sequencing of 685 genes or ThunderBolt Myeloid Panel (RainDance Technologies), a NGS panel covering 49 genes frequently mutated in myeloid disorders, as described previously^42^. CALGB/Alliance samples were sequenced using a NGS panel covering 80 cancer and/or leukemia associated genes as described previously^43^. Patient samples were selected based on the presence of *NPM1* mutations with additional co-occurring mutations of *DNMT3A, TET2, IDH1/2, NRAS,* and/or *FLT3* due to their high frequencies in AML patients. For longitudinal samples, only diagnosis samples were molecularly profiled. Patients for longitudinal analysis were prioritized if they had *TET2* or *IDH2* co-occurring mutations at diagnosis. Treatment information for patients with longitudinal samples is summarized in Extended Table 2 and displayed in Extended Data Figure 1.

### Single-cell DNA and protein (scDNA+Protein) library preparation and sequencing

Patient samples were thawed, washed with FACS buffer, filtered into single cell suspensions, and quantified using a CellDrop (Denovix). Cells (1x10^6^ viable cells) were then incubated with TruStainFcX and Tapestri blocking buffer for 15min on ice followed by a 30min incubation with the TotalSeqD Antibody Cocktail on ice. A select number of samples were also supplemented with 2μL of TotalSeqD CD135 during this step. Stained cells were then washed three times with Cell Staining Buffer. Cells were filtered through a Flowmi cell strainer (vendor), centrifuged, resuspended with Tapestri cell buffer, quantified and loaded into the Tapestri microfluidics cartridge. Single cells were encapsulated, lysed, and barcoded as described previously^8^. DNA PCR products and Protein products were isolated and purified using AMpure XP beads and Streptavidin C1 beads, respectively. DNA PCR products and C1-bead immobilized Protein products were each used as PCR templates for DNA and Protein-derived DNA library generations, respectively followed by a final purification using AMpure XP beads. DNA and Protein derived libraries were quantified using an Agilent Bioanalyzer and Qubit (Invitrogen) and pooled for sequencing on an Illumina NovaSeq6000. Sequencing of pooled libraries were performed by the MSKCC Integrated Genomics Core and the DNA Genomic Sequencing shared facility at CCHMC. scDNA+Protein sequencing metrics for all samples are provided in Extended Table 1.

### CITE-seq

Patient samples were thawed, washed, and quantified as above. Cells were then stained with 7-AAD (Biolegend) and viable cells (200,000 per sample) sorted using a Sony MA900 cell sorter. A previously used^44^ custom Total-seq A oligo-conjugated antibody panel from Biolegend was used to stain live sorted AML cells. Sorted cells (200,000/sample) were stained for 60 minutes on ice, washed using laminar flow (Curiox), and resuspended prior to counting. Cells (16,000 per well) were loaded using 10X Chromium Gene Expression 3’ version 3.1 kit (1000268, 10X Genomics). Emulsion, GEM collection, clean-up and cDNA amplification were performed according to 10X Genomic protocol. Library preparation was performed according to the manufacturer’s protocols. Final transcriptome libraries were quantified and analyzed using a Qubit dsDNA HS assay kit (Q32854, Invitrogen), a High-Sensitivity DNA kit (5067-4626, Agilent Technologies) on a 2100 Bioanalyzer (G2939BA, Agilent Technologies) and a KAPA HiFi library quantification kit (KK4824, Roche). Dual-indexed transcriptome libraries were pooled and sequenced on two X plus lanes with the PE100 settings (Illumina). BCL files were demultiplexed into fastq files for CellRanger V7.1.2 input. The transcriptome was mapped to hg38 reference genomes for downstream analysis and visualization.

### CITE-seq analysis

All Cell Ranger-produced count matrices underwent ambient RNA exclusion using the software SoupX^45^ with a contamination fraction of 15% and quality control filtering by HTO and Seurat V4^46^. Ambient corrected transcriptome counts and associated ADT counts were supplied as input to the software TotalVI to obtain normalized and denoised ADT counts. To derive clusters from our previously published human bone marrow progenitor atlas^25^, the software cellHarmony^47^ was used to transfer labels from CPTT normalized expression centroids from synapse. Cell annotations from our previously generated human bone marrow CITE-seq atlas were projected onto the merged dataset using cellHarmony. Cells with a poor mapping score to the final clusters (linear support vector classification coefficient > 0) were excluded from the analysis (for example, doublets). Differential gene expression analysis was performed between these three samples with cellHarmony at a threshold of log2 fold change >1.2 and *P* value <0.05.

### Single cell DNA sequencing analysis

Sequencing reads were trimmed, aligned to the human genome (hg19), assigned barcodes, and genotyped were called with GATK by the cloud-based Mission Bio Tapestri v2 pipeline. Processed H5 files were further analyzed using the scDNA package (https://github.com/bowmanr/scDNA, v1.01) in R v4.3. In the scDNA package, H5 files from the Mission Bio Tapestri pipeline were used as input and variants of interest were identified in the following genes *DNMT3A*, *TET2*, *IDH1*, *IDH2*, *NPM1*, *FLT3*, *PTPN11*, *NRAS*, and *KRAS*. All variants included in this study were manually investigated in IGV. We selected exonic, non-synonymous variants that were genotyped in >50% of cells assayed and had a computed VAF >1%. For samples acquired at remission we decreased the VAF cutoff to 0.1%. We further refined the variant list to exclude those that were either 1) confirmed SNPs, 2) were recurrently mutated at a fixed VAF broadly across the cohort, 3) only represented in low quality reads or clipped reads visual inspection in IGV. Excluded variants included: TET2.I1762V, TET2.961*, TET2.Y1579*, TET2.A1283T, TET2.L1721W, DNMT3A.F772C, NRAS.L56P, NRAS.T58A, NRAS.L56Q, NRAS.D57N, DNMT3A.I310S, NRAS.T58I, DNMT3A.I292S, DNMT3A.L888Q, DNMT3A.L888P, PTPN11.L525R, DNMT3A.N489T, DNMT3A.K429T, DNMT3A.N757T, NRAS.T58P, NRAS.D57Y, NRAS.Q61P, TET2.Q618H, TET2.L1819F, DNMT3A.Y481S, FLT3.N847T, TET2.A584T, TET2.A584P, DNMT3A.F290L. In the case of paired samples, we included variants that were below the 1% VAF threshold if they were present in another sample in the pair so as to identify rare subclonal events. Following variant selection, the ‘tapestri_h5_to_sce’ function from the scDNA package was used to generate a SingleCellExperiment class object using the default cutoffs of depth (DP) >10, genotype quality (GQ) >30, and allele frequency (AF) variance >25. The AF variance refers to the maximum deviation from 50% by which a heterozygous call from GATK should be masked as inaccurate. Finally, we retained variants that passed all three of these filters in over 80% of cells. Only cells that passed all three filters were included in the final analysis and were termed “Complete” cells, indicating they received a reliable genotype for all genes of interest. Following variant identification, clones were identified and statistically summarized using the ‘enumerate_clones’ and ‘compute_clone_statistics’ functions respectively.

### Single cell DNA+Protein (scDNA+Protein) sequencing analysis

Following genotyping and clone enumeration above, protein matrices were extracted from H5 files from the tapestry pipeline using the scDNA package. The SingleCellExperiment object was converted to a Seurat object (v5.1) and genotype information was stored as metadata. For global protein analyses across all samples, all complete cells identified above were bound to a single protein matrix, and each sample was downsampled to 7,000 cells. For samples with <7,000 cells, all cells were included. Protein data was normalized across cells using CLR (margin=2), scaled across all samples, and analyzed by PCA. Samples were integrated with Harmony, then clustered (SLM) and visualized by UMAP^48^. Clusters with high protein counts, high protein feature abundance (e.g. possessed every antibody) and high abundance of IgG antibodies were considered ‘dead’ and removed from the analysis. Following dead cells removal, we reran the steps above from Normalization through to UMAPs. A similar process was undertaken for patient sample pairs, starting from a raw read count matrix that only contained the patient sample of interest. Cell type calls were performed by manual interpretation of protein expression. Data was visualized using Seurat, ggplot2 and scCustomize packages (https://samuel-marsh.github.io/scCustomize/).

### Statistical analysis

Comparisons of clonal architecture metrics were analyzed by Kruskal-Wallis tests. Two-way ANOVA tests were used to analyze clonal synergies between co-mutations. A Wilcoxon Rank Test was used to assess significant differences in protein expression in the scDNA+Protein data.

### Plotting and graphical representations

Clonal architecture metric plots (Fig. 2, Extended Data Fig. 2), clonal frequency plots (Fig. 3-4, Extended Data Fig. 3-4), and treatment response courses (Extended Data Fig. 1) were generated using GraphPad Prism. Error bars depict standard error of the mean. The oncoprint in Fig. 1a was generated in R using oncoPrint package. For patients who had more than one sample in the cohort (n=8), we only included one sample prioritizing the diagnosis sample, if possible. No complete response (CR) samples were included in the oncoprint. UMAP data was plotted using the ggplot2 package in R. Other data processing was performed in R utilizing packages including: tidyr, dplyr, RColorbrewer, pals, and cowplot. Differentially expressed genes and ADT counts (Fig. 6) were plotted in heatmaps by Alt Analyze^49^ and used to identify common perturbed biological processes. GO Biological Processes were plotted based on Z-score and adjusted P values (Fig. 6e). The values in each row were normalized to the median of the row and used to derive the heatmaps. Network graph in Figure 6f was plotted using Cytoscape^50^ with log fold gene expression denoted by color of circles (high = red, low = blue).

## Data availability

All scripts and processed data files are available for DNA+Protein analyses at https://github.com/bowmanr/scDNA. Raw data files are available upon request from the authors and are being uploaded to dbGAP prior to final publication.

## Code availability

Once processed through the Tapestri pipeline, samples were initially filtered and analyzed using a custom code scripted in R (github.com/bowmanr/scDNA). Scripts for CITE-seq processing through Seurat can be found at https://github.com/satijalab/seurat. The AltAnalyze v.2.1.4 graphical user interface was utilized for the cellHarmony and differential expression analyses as described. GraphPad Prism v.10 was used for sample and cell frequency plotting.

## Acknowledgements

We acknowledge the use of the CCHMC Genomics Sequencing Core (RRID:SCR_022630) and the MSKCC Integrated Genomics Core (supported by NIH P30 CA008748) for library sequencing. M.D. is supported by a CCHMC Strauss Clinical Fellow Award and an NIH training grant (T32 CA236764-5). B.K. is supported by a Leukemia & Lymphoma Society (LLS) Career Development Fellow Award. L.A.M. is supported by a National Cancer Institute grant (R00 CA252005) and an American Society of Hematology (ASH) Junior Faculty Scholar award. R.L.B. is supported by a National Cancer Institute grant (R00 CA248460) and an ASH Junior Faculty Scholar award. This work was also supported by the ASH Junior Faculty Scholar award and the NCI R00 award to L.A.M. The authors are grateful to the patients who consented to participate in these clinical trials and the families who supported them; to Christopher Manring and the CALGB/Alliance Leukemia Tissue Bank at The Ohio State University Comprehensive Cancer Center, Columbus, OH for sample processing and storage services; and to Lisa J. Sterling for data management.

## Author contributions

L.A.M., R.L.B., R.L.L., H.L.G conceptualized studies. R.L.B., and M.B. designed and optimized single cell DNA/Protein experimental methodologies and bioinformatic workflow. X.Z., B.K., N.S., and H.L.G. designed and optimized CITE-seq protocol and the bioinformatic workflow. D.N., R.S., K.M., A.J.C., A.K.E., R.L.L., and J.C.B. provided de-identified patient samples and annotated clinical information. M.D., D.L., B.K., X.Z., and L.A.M. performed library preparation and sequencing. X.Z. M.B., N.S., and R.L.B. performed all computational multiomic analysis. M.D., D.L., X.Z., Z.W., R.L.B. and L.A.M. generated manuscript figures. L.A.M. funded the study. M.D., D.L., and L.A.M. wrote and edited the manuscript with contributing edits from K.M., D.S., A.K.E., H.L.G., and R.L.B.. All authors read the manuscript and agreed on the final version.

